# APDeeM: A machine Learning strategy towards Effective Peptide Vaccine Candidates Identification against Different Types of Viruses

**DOI:** 10.1101/2025.08.25.671769

**Authors:** Mohammad Uzzal Hossain, Md. Romzan Alom, SM Sajid Hasan, Mohammad Nazmus Sakib, Marjia Akter Suchi, Zeba Sanjida, A.B.Z. Naimur Rahman, Arittra Bhattacharjee, Zeshan Mahmud Chowdhury, Ishtiaque Ahammad, Muhammad Aminur Rahman, Saiful Azad, Md. Salimullah

## Abstract

Viral infections pose significant global health challenges, underscoring the urgent need for improved medications. Nevertheless, traditional medicinal approaches depend significantly on labor-intensive laboratory tests, which impede efficient identification and prolong vaccine development, particularly when screening a huge number of samples. To address these obstacles, we present a comprehensive Antiviral Peptide (AVP) Detection Dataset, comprising 14 unique features to improve the characterization of antiviral and non-antiviral peptides. Subsequently, we introduce the Antiviral Peptide detection enhanced by Ensemble Machine Learning (APDeeM) system. This advanced computational framework considerably reduces the time required for AVP detection by utilizing ensemble learning methodologies. The APDeeM system incorporates Gradient Boosting, Random Forest, K-Nearest Neighbors (KNN), and AdaBoost algorithms to facilitate the swift selection of AVP candidates without requiring urgent laboratory testing. Our proposed ensemble methodology showed superior performance, with an accuracy of 85.99%, F1 score of 87.60%, recall of 88.91%, and precision of 86.32%, exceeding the efficacy of all tested antiviral peptide prediction models in this research. The APDeeM approach signifies a substantial improvement over conventional detection techniques, expediting the identification of prospective vaccine candidates and facilitating the advancement of more effective antiviral peptides. The most promising AVP candidates may urge laboratory validation, optimize resources, and accelerate vaccine development.

## Introduction

Viral infections pose a major threat to global public health by impacting morbidity, mortality, and socio-economic conditions^1^. Various viral outbreaks, notably SARS-CoV-2, Influenza, and Human Immunodeficiency Virus (HIV), outlined the urgency of effective antiviral treatments and/or vaccines ^2,3^. For this reason, several approaches have been developed including both antiviral drugs and vaccines to fight against viral infections ^4,5^. However, the manifestation of viral strains rapidly, contrary to the slow and extensive development process of vaccines, has limited the beneficial impacts of these antiviral strategies ^6,7^. Hence, it has become very important to search for innovative treatment methods against viral activities that can provide support rapidly and efficiently.

One of the innovative treatments that can work alternatively to conventional approaches is antiviral peptides (AVPs) ^8^. These are short in sequence and made of amino acid chains. Distinct antiviral properties make them unique to combat against a diverse range of viral entities. These properties may include, but are not limited to, blocking the entry of the virus into host cells, potentially halting viral replication, and boosting the immunological activities within the host ^9,10^. Nevertheless, what makes them attractive is their broad-spectrum activity, low risk in generating resistance, and rapid action upon viral infections ^11,12^. Although they possess advantages over traditional antiviral treatments, identifying and characterizing these AVPs remains a very challenging task ^13^. Generally, laboratory experiments are utilized in discovering the AVPs, but the heavy cost and the long duration of the processes are deemed unrealistic when it is necessary to screen peptide libraries on a large scale ^14,15^.

Recent developments in machine learning (ML) and deep learning (DL) can computationally advance our search for potential AVPs ^16^. In this way, algorithms targeted for this purpose can leverage computational resources for enhanced identification of complexity in diverse peptide sequences and derive specific patterns for AVPs. Moreover, bigger datasets can be explored with great ease and over-reliance on experimental tools can be decreased ^17^. Realizing these benefits of in-silico approaches does not appear to be modifying the subject computational models for AVP predictions. But these models often suffer from various limitations, such as vulnerability to the imbalance in data, inconsistency in overall performance (specifically in various preprocessing steps), and the usage of minimal feature sets ^14,18^. Therefore, these shortcomings hinder the development of sustainable AVP detection methods.

Amid the presence of shortcomings in the domain of machine learning intended for AVP prediction, several models tried to address major challenges, including DeepAVP ^19^, Deep-AVPiden ^20^, Deep-AVPpred ^21^, and AntiCVP-Deep ^22^. In DeepAVP, integrated both LSTM and CNN, categorizing it as a hybrid model. Although it was initially promising, leveraging a small set of features greatly reduces its generalizability. On the other hand, Deep-AVPiden, achieving high recall values was shadowed by low precision and accuracy, which impeded its capability to utilize imbalanced datasets. Another model, Deep-AVPpred was inspired by ResNet and performs moderately; however, its reliability in AVP identification is compromised due to the lack of robustness. Furthermore, AntiCVP-Deep integrated BiLSTM and self-attention for strong predictive performance, but its generalizability is limited by synthetic data and model complexity.

Employing a genetic algorithm was another important approach that was introduced by the Ensemble Learner model ^23^. It achieved a high AUC score but dropped in F1 score and accuracy. Hence, it demonstrated trade-offs where increasing complexity decreased performance. Furthermore, models built on SVM and Random Forest often face difficulties due to imbalances in datasets and, therefore, become unable to grasp the distinct properties of AVPs, particularly physicochemical and immunological ones ^24,25^. These failures strongly necessitate the importance of frameworks focusing on the comprehensive prediction of AVPs.

In search of comprehensive frameworks, we present a novel computational framework titled APDeeM that identifies antiviral peptides. This framework integrates datasets with 14 distinct features, including net charge, molecular weight hydrophobicity, and the Boman index for capturing the peptide-viral component interactions. Additionally, it uses advanced preprocessing techniques and a machine learning model based on ensemble learning to accurately and reliably predict AVPs. Further, many algorithms such as Gradient Boosting, Random Forest, K-Nearest Neighbors (KNN), and AdaBoost have been combined, which upgrades the performance level compared with similar models. Therefore, it could accelerate the detection of potential AVPs and demonstrate its ability in the search for antiviral therapies.

### Data Preparation and Methodology

In this research, **Figure 1** illustrates the proposed methodology of antiviral peptide detection enhanced by Ensemble Machine Learning (APDeeM) system. The workflow begins with dataset preparation and advanced preprocessing, followed by model training using the designed ensemble algorithms, and concludes with performance evaluation using benchmark metrics, ensuring the system’s effectiveness in identifying potential antiviral peptides.

**Figure 1.**
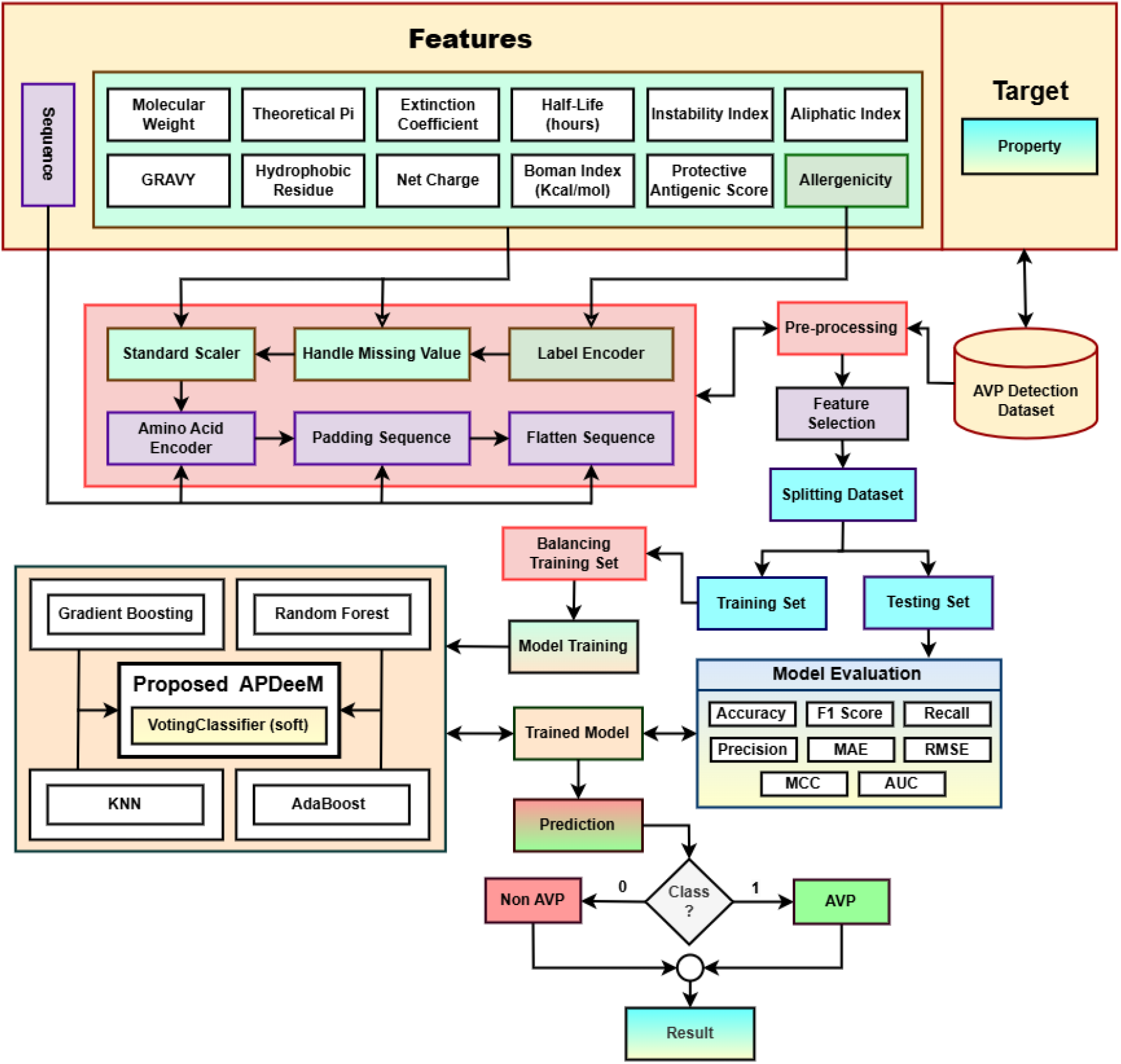
Proposed methodology of the APDeeM system for antiviral peptide prediction pipeline. AVP Detection dataset: shows the attributes of the entire dataset that contain peptide sequenc characteristics. Data preprocessing: illustrates several preprocessing tasks done on different attributes of the created dataset. Model training: the balanced training set of data is learned using various machin learning, deep learning, and our constructed ensemble models. Model evaluation: shows the evaluation metrics to select the best model for the target prediction.

### Dataset Description

In this study, we created a new dataset that contains 3,886 entries, each representing biological sequences with various molecular and immunological properties. We explored a variety of computational web tools using those peptide sequences to gather key properties associated with peptide characteristics in order to gain a greater understanding of the model.

We obtained the Theoretical pi, Extinction Coefficient, Grand Average of Hydropathy (GRAVY), Molecular Weight, Half-life, Instability Index, and Aliphatic Index using “Protein Identification and Analysis Tools” from Expasy ^26^. The Hydrophobic Residues, Net Charge, and Bomen Index were acquired using “APD3 Tool” ^27^. Additionally, Protective Antigenic Score and Allergenicity were picked up using “VaxiJen Tool” ^28^ and “AllerCatPro 2.0” ^29^, respectively. Some of our authors manually annotated these attribute values to ensure a comprehensive dataset.

The data was classified into two distinct classes. Property value of 1 means AVP (Antiviral Peptide), which has 2082 samples, and 0 means non-AVP, which has 1804 samples (**Figure 2**).

**Figure 2.**
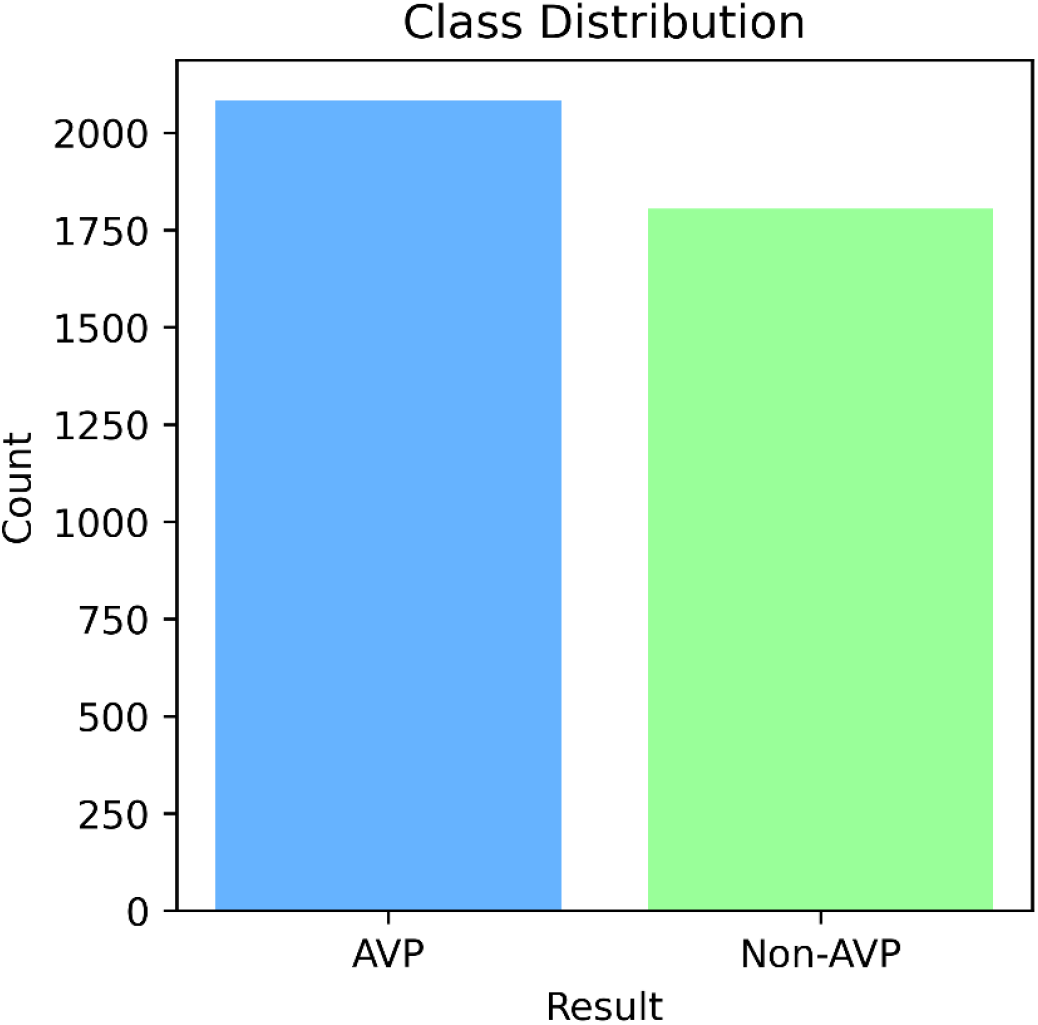
Class distribution of the property class. The blue line indicates the distribution of antiviral peptides (AVP), with a count of 2082, while the green line specifies the non-antiviral count, which is 1804 in total.

### Data Pre-processing

This study employed the Antiviral Peptide (AVP) Detection Dataset, which posed several challenges such as diverse categorical features, missing values, and significant variability in numerical data. To enhance the reliability and uniformity of the dataset, a comprehensive set of preprocessing techniques was applied, as detailed below:

### Label Encoding

Label encoding ^30^ transforms categorical data into numerical values by assigning a unique integer to each category using **Equation 1**.

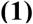

Where *L(x)* is the label-encoded value for category *x, C* is the set of unique categories, and Index*(x, C)* is the integer index based on category order.

### Missing Value Handling

The preprocessing involved replacing missing values with NaN, then filling categorical missing values with the mode using **Equation 2**.

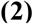

Where *x*_*i*_ represents the value of the *i-th* position in the attribute, and *m* denotes the mode of the column.

The numerical missing values were imputed with the mean of its respective attributes using

### Equation 3

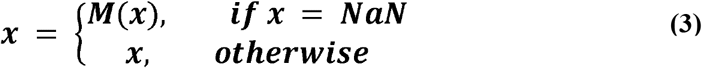

Where *M* denotes the mean value of the column.

### Standard-Scaler

After converting attributes to numerical values, StandardScaler from scikit-learn ^31^ was applied to standardize the distribution, resulting in a mean of 0 and a standard deviation of 1 using **Equation 4**.

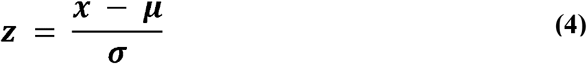

Where *z* is the scaled value, *x* is the original value, μ is the mean, and σ is the standard deviation.

### Encoding Amino Acid Sequences

Each amino acid was mapped to a unique index, with the encoded sequence *S*′ represented by

### Equation 5

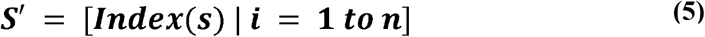

Where *S*_*i*_ is the *i*-th amino acid, Index(*S*_*i*_) is its assigned index, and *n* is the sequence length.

### Padding Sequences

To ensure that all sequences were of equal length, they were padded to a fixed length *L*_*max*_ using zero padding, represented by **Equation 6**.

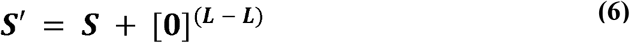

Where *S*′ is the padded sequence, *S* is the original sequence, *L* is the original length, and *L*_*max*_ is the desired fixed length.

### Combining the Flattened Sequence Data with Other Features

Afterwards, the encoded and padded sequences were flattened into a 2D structure and combined with additional features to form the processed data *X*_*fina*_ using **Equation 7**.

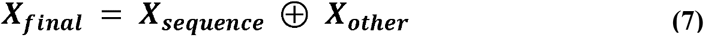

Where *X*_*sequence*_ is the flattened sequence data, and *X*_*other*_ represents the additional features.

### Feature Selection

The Pearson correlation coefficient matrix was used to show linear relationships between variables ^32^ (**Figure 3**).

**Figure 3.**
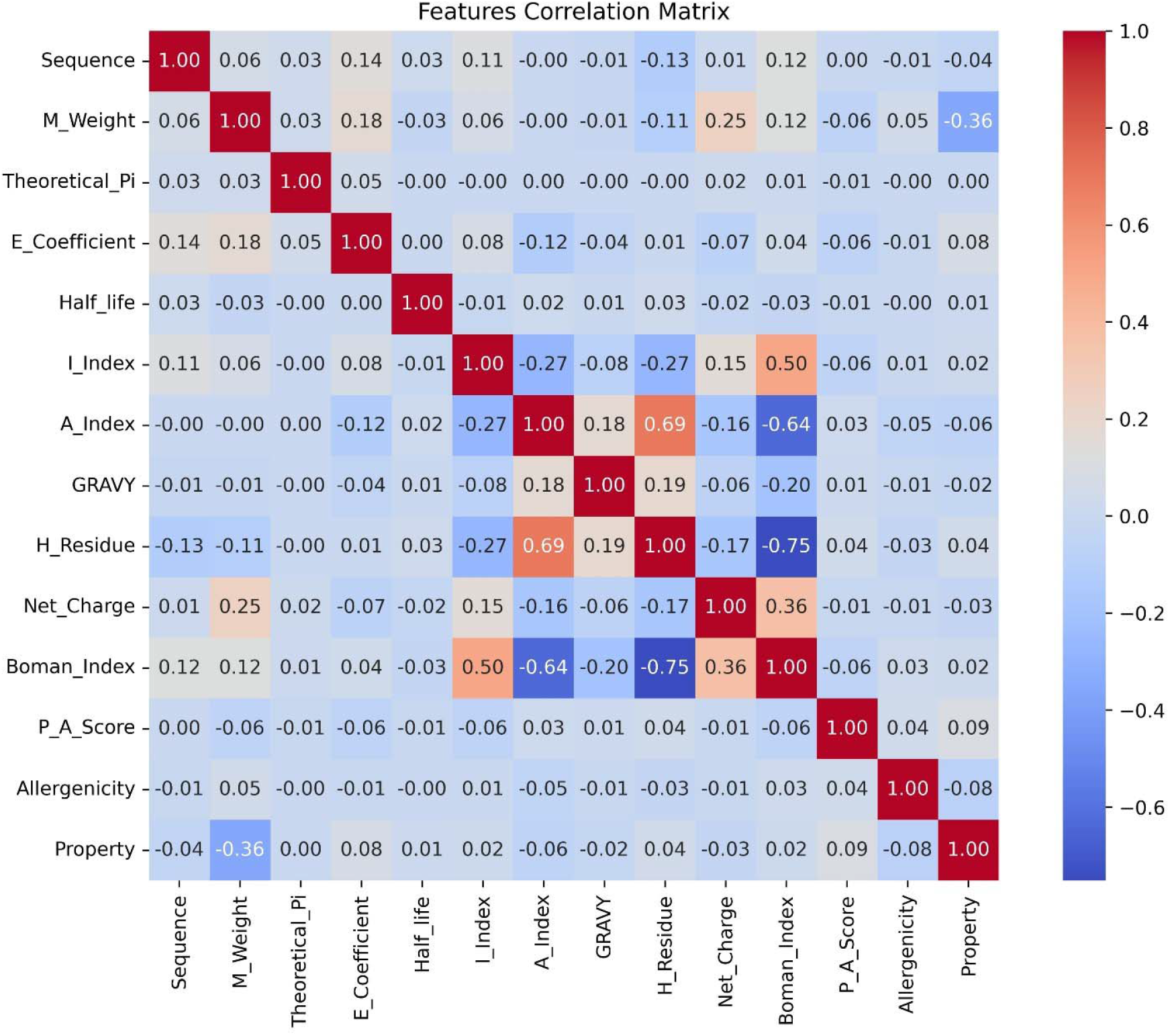
Correlation matrix for different features. The hydrophobic residue (H Residue) and Boman index had a highly negative relationship (Pearson correlation coefficient, r = −0.75), followed by the aliphatic index (A Index) and Boman index (r = −0.64). In contrast, the aliphatic index (A index) and hydrophobic residue (H Residue) had a high positive correlation (r = 0.69), and subsequently, the Boman index and instability index (I Index) had a good positive relationship (r = 0.5).

Correlation values range from −1 (strong negative) to 1 (strong positive), with 0 indicating no correlation.

### Splitting Dataset

The entire dataset was divided into two distinct sets, the training set and the testing set, using the train test split method from the scikit-learn ^33^ with an 80:20 ratio. The training set consisted of 3108 samples (1649 AVP and 1459 non-AVP), while the testing set contained 778 samples (433 AVP and 347 non-AVP).

### SMOTE for Data Balancing

Synthetic Minority Oversampling Technique (SMOTE) was applied to the training data to keep the test set unbiased, generating synthetic minority samples instead of duplicates ^34^. After balancing the train set, the training set contained 3298 samples with an equal number of AVP and non-AVP instances (**Figure 4**).

**Figure 4.**
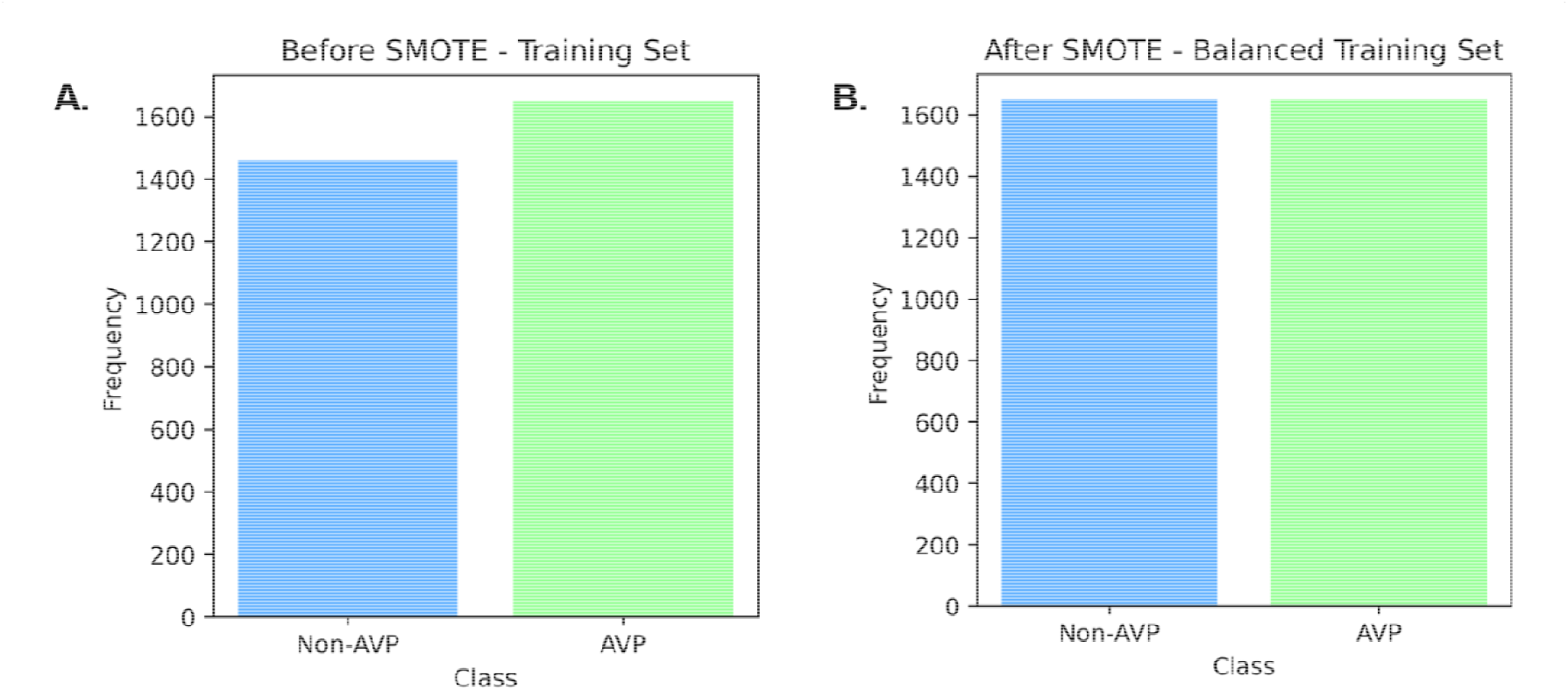
Class distribution on training set of the AVPD dataset. (A) Before SMOTE is applied on the training set (1649 AVP and 1459 non-AVP sample) (B) After SMOTE is applied on the training set (1649 AVP and 1649 non-AVP sample).

### Model Development and Training

In this research, we trained a comprehensive set of 21 models, incorporating both traditional machine learning and deep learning techniques, alongside our proposed ensemble models. This diverse array of models was applied to our dataset to facilitate equitable model training. The traditional machine learning models included Support Vector Machine (SVM), Decision Tree, Extra Tree (ET), Random Forest, Logistic Regression, Gradient Boosting, K-Nearest Neighbor (KNN), Neural Network, Naïve Bayes, AdaBoost, XGBoost, and Balanced Random Forest ^24,35–37^.

Additionally, we integrated five notable prior works in this field: the DeepAVP model (LSTM+CNN channels) ^19^, Deep-AVPiden (temporal convolutional networks) ^20^, the Deep-AVPpred model (ResNet-1D) ^21^, the AntiCVP-Deep model (bidirectional LSTM with attention mechanisms) ^22^, and the Ensemble Learner model (combining XGBoost+ KNN+ET+SVM+AdaBoost with genetic algorithms) ^23^.

During the model creation process, two ensemble learning techniques were implemented in our research. Soft voting was applied to constructing the Ensemble 1 and Ensemble 3 models, whereas Ensemble 2 and 4 used stacking of the classifiers. Both soft voting and stacking of the classifiers combine multiple base classifiers to enhance predictive performance.

**Ensemble 1:** This ensemble combined four machine learning algorithms (Gradient Boosting, Random Forest, K-Nearest Neighbors (KNN), and AdaBoost) that employed a voting classifier to aggregate the predictions from these diverse algorithms to enhance predictive performance. The final prediction was obtained by averaging the predicted probabilities from each individual classifier.

Let *x* be an input sample, *C* be the number of possible classes, *M* be the number of base classifiers which is 4, *P*_*m*_ *(y*_*c*_ | *x)* be the probability predicted by the *m*-th classifier for class *y*_*c*_, and *P (y*_*c*_ | *x)* be the final predicted probability for class *y*_*c*_. The soft voting classifier computes the final probability for class *y*_*c*_ (**Equation 8**).

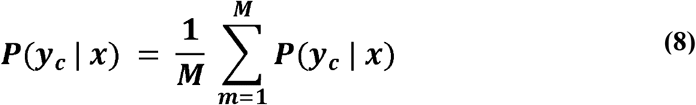

The final prediction class ŷ is determined by the **Equation 9**.

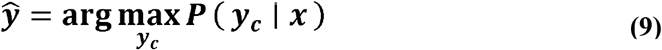

Where *P (y*_*c*_ | *x)* represents the average class probability for class *y*_*c*_.

This probability is computed as the average of the individual model probabilities using **Equation 10**.

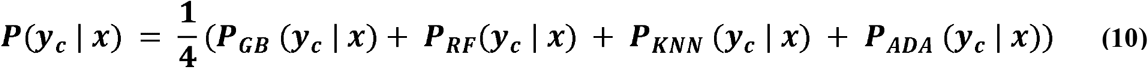

Where *P*_GB_*(yc* | *x), P*_RF_*(yc* | *x), P*_KNN_*(yc* | *x)*, and *P*_ADA_*(yc* | *x)* are the individual class probabilities predicted by each model (Gradient Boosting, Random Forest, K-Nearest Neighbors, AdaBoost).

From the classifiers, Gradient Boosting constructed a group of 200 decision trees, each with a maximum depth of 10 and being weighted by a constant learning rate 0.1 (**Equation 11**).

Likewise, the Random Forest model used 200 trees, each built to a maximum depth of 10, and the trees were constructed using bootstrap samples, requiring at least one sample per leaf and a minimum of two samples for splitting a node (**Equation 12**). Using the Minkowski distance, the KNN model used the five nearest training samples to predict the probability class *y*_*c*_ based on their class distribution (**Equation 13**). The final classifier AdaBoost used 200 base learners, each with a weight of 0.1, and like the gradient boosting model, it sequentially added those weak learners to build a progressive model (**Equation 14**).

Gradient Boosting (GB):

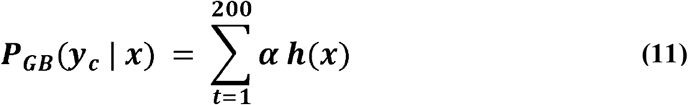

Random Forest (RF):

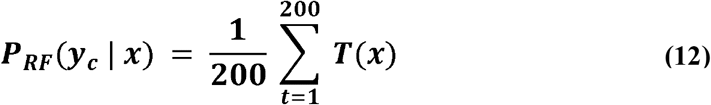

K-Nearest Neighbors (KNN):

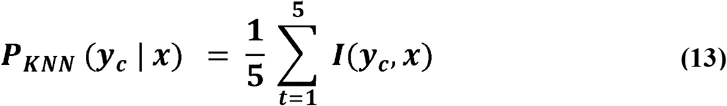

AdaBoost(ADA):

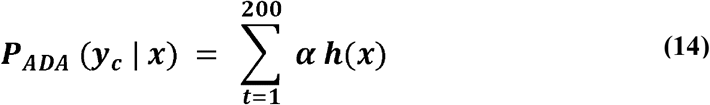

**Ensemble 2:** A stacking classifier with Gradient Boosting, K-Nearest Neighbors, and AdaBoost were utilized as base models of this ensemble model and Random Forest as the meta-estimator. This approach capitalized on the strengths of each base model, while the Random Forest final estimator synthesized their outputs to improve generalization.

**Ensemble 3:** This is the third ensemble model that incorporated XGBoost, Balanced Random Forest, K-Nearest Neighbors, and AdaBoost in a voting classifier. By combining these models through voting, Ensemble 3 achieved balanced performance, improved minority class detection and overall accuracy.

**Ensemble 4:** The Ensemble 4 followed a stacking configuration with XGBoost, K-Nearest Neighbors, and AdaBoost as base models, and Balanced Random Forest as the final estimator to handle imbalanced data effectively.

### Model Evaluation and Prediction

After completing the training process, all trained models were evaluated using a designated test set. Various metrics were employed to assess their performance, including Accuracy, F1 Score, Recall, Precision, AUC Score, Matthews Correlation Coefficient (MCC), Mean Absolute Error (MAE), and Root Mean Square Error (RMSE) ^38^. Based on the evaluation results, the best performing model was selected for predicting antiviral peptides.

### Result Analysis Experimental Setup

The study was conducted on the Kaggle platform utilizing its default hardware configuration, which includes an Intel Xeon CPU, 13GB of RAM, and approximately 80 GB of disk space-suitable for both machine learning and deep learning tasks.

The environment was built using Python 3.7, with libraries including scikit-learn, pandas, numpy, seaborn, imblearn and matplotlib, and TensorFlow supporting data preprocessing, model development, visualization, and evaluation ^39^. A wide range of hyperparameters was employed to control the learning dynamics and boost the performance of the 21 trained models. The selected hyperparameters was based on iterative experimentation and validation (see **Supplementary Table 1, Table 2**, and **Table 3**). Optimal configurations contributed to the robustness and reliability of the predictive models.

### Performance Measurement Parameters

In this study, model performance was evaluated using the confusion matrix, which summarizes predictions by counting true positives (TP), true negatives (TN), false positives (FP), and false negatives (FN) ^40^. These values were used to compute key evaluation metrics, offering insights into the models’ predictive effectiveness. The following metrics were employed for assessment:

- **Accuracy (%):** Indicates the proportion of correctly classified instances, calculated using**Equation 15**.

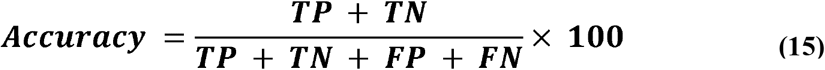
- **F1 Score (%):** Represents the harmonic mean of precision and recall, computed using**Equation 16**.

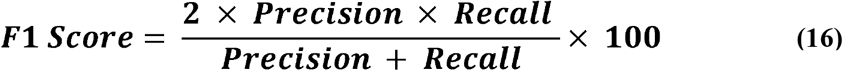
- **Recall (%):** Indicates the proportion of actual positives correctly identified by the model, calculated using **Equation 17**.

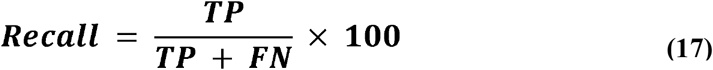
- **Precision (%):** Indicates the proportion of positive predictions that are actually correct, determined using **Equation 18**.

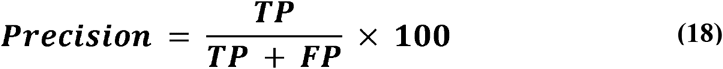
- **AUC (%):** Measures the model’s class-separating ability, evaluated using **Equation 19**.

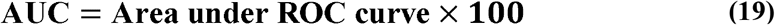
- **MCC (%):** Measures balanced performance using all confusion matrix elements, computed using **Equation 20**.

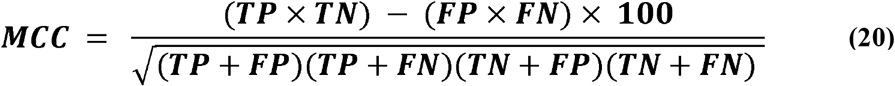
- **MAE (%):** Quantifies the average absolute difference between predictions and actual values, calculated using **Equation 21**.

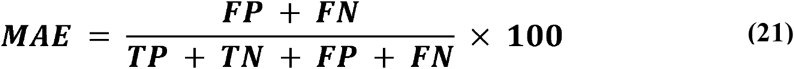
- **RMSE (%):** Measures the square root of the average squared differences between predicted and actual values, determined using **Equation 22**.

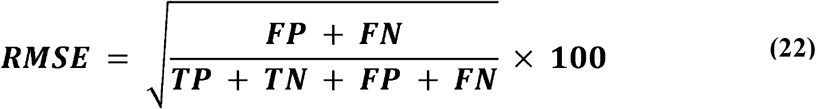

### Comparative Analysis

Is this study, **Table 1** and **Supplementary Figures 1–21** present the confusion matrix values for all trained models.

**Table 1:**
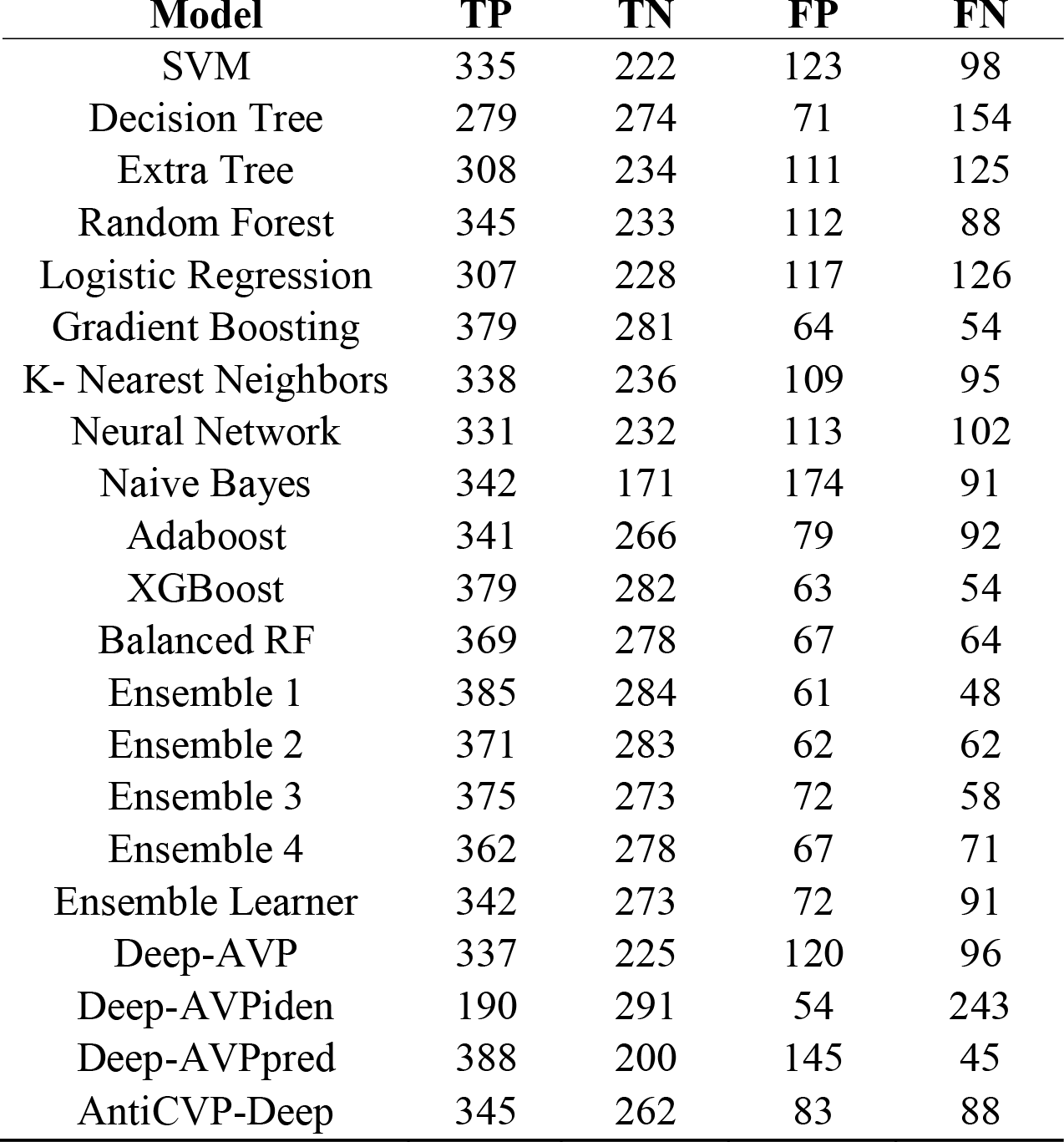
Confusion matrix summary exhibiting the true positive (TP), true negative (TN), false positive (FP), and false negative (FN) for each model.

| **Model** | **TP** | **TN** | **FP** | **FN** |
| --- | --- | --- | --- | --- |
| SVM | 335 | 222 | 123 | 98 |
| Decision Tree | 279 | 274 | 71 | 154 |
| Extra Tree | 308 | 234 | 111 | 125 |
| Random Forest | 345 | 233 | 112 | 88 |
| Logistic Regression | 307 | 228 | 117 | 126 |
| Gradient Boosting | 379 | 281 | 64 | 54 |
| K- Nearest Neighbors | 338 | 236 | 109 | 95 |
| Neural Network | 331 | 232 | 113 | 102 |
| Naive Bayes | 342 | 171 | 174 | 91 |
| Adaboost | 341 | 266 | 79 | 92 |
| XGBoost | 379 | 282 | 63 | 54 |
| Balanced RF | 369 | 278 | 67 | 64 |
| Ensemble 1 | 385 | 284 | 61 | 48 |
| Ensemble 2 | 371 | 283 | 62 | 62 |
| Ensemble 3 | 375 | 273 | 72 | 58 |
| Ensemble 4 | 362 | 278 | 67 | 71 |
| Ensemble Learner | 342 | 273 | 72 | 91 |
| Deep-AVP | 337 | 225 | 120 | 96 |
| Deep-AVPiden | 190 | 291 | 54 | 243 |
| Deep-AVPpred | 388 | 200 | 145 | 45 |
| AntiCVP-Deep | 345 | 262 | 83 | 88 |

In this investigation, (**Supplementary Table 4**) presents a comprehensive comparison of all trained models across multiple performance metrics using **Table 1**. Our proposed Ensemble 1 model demonstrated superior performance, achieving the highest accuracy, F1 score, recall, precision, and MCC, with respective values of 85.99% (**Figure 5**), 87.6% (**Figure 6**), 88.91% (**Figure 7**), 86.32% (**Figure 8**), and 71.55% (**Figure 10**).

**Figure 5.**
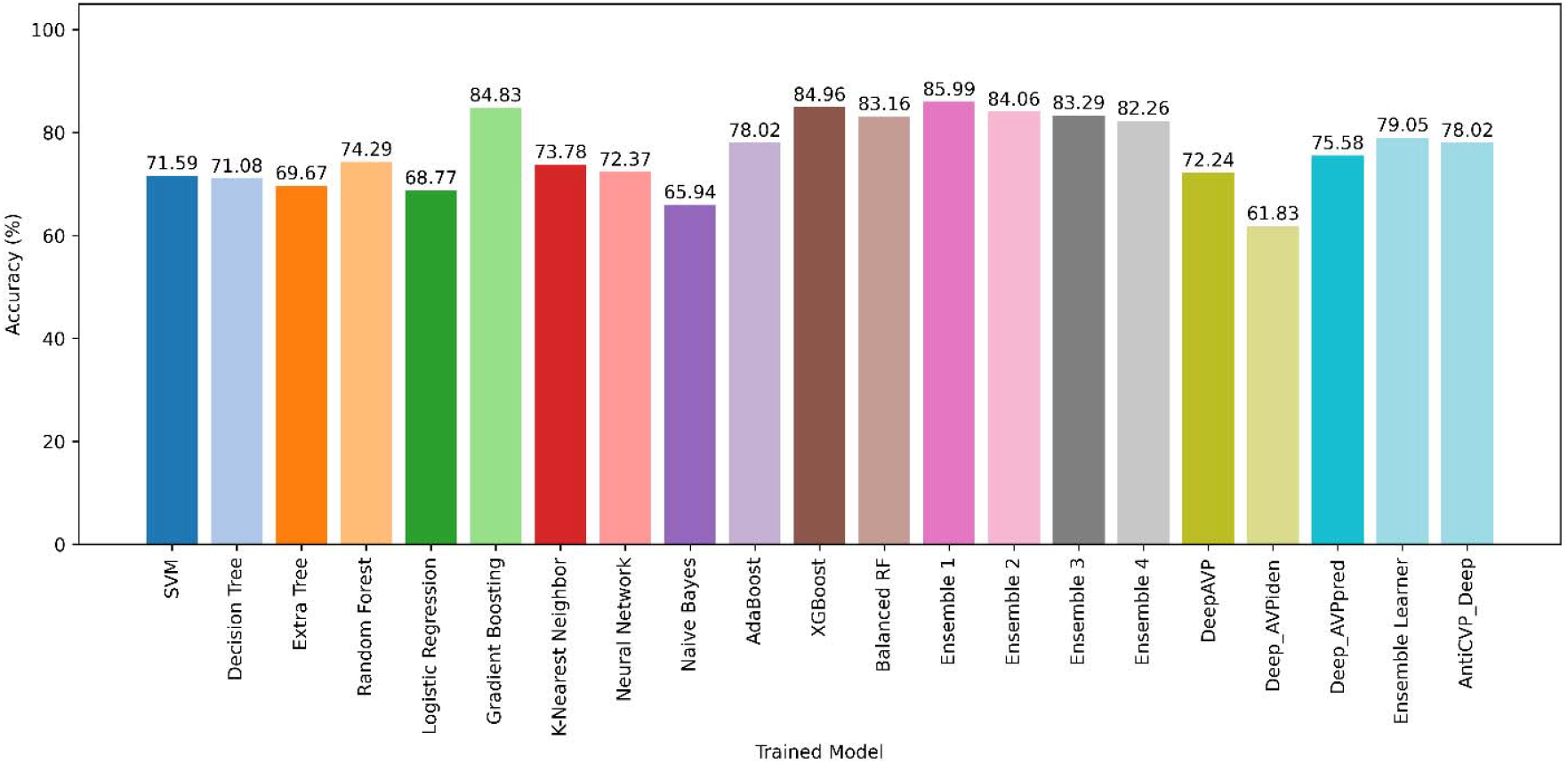
Accuracy comparison of the models. The performance of each model in terms of accuracy is demonstrated here. The best accuracy is achieved from ensemble model 1 (Gradient Boosting, Random Forest, KNN, AdaBoost). With an accuracy of 85.99%, the dark pink line represents the Ensemble 1 model.

**Figure 6.**
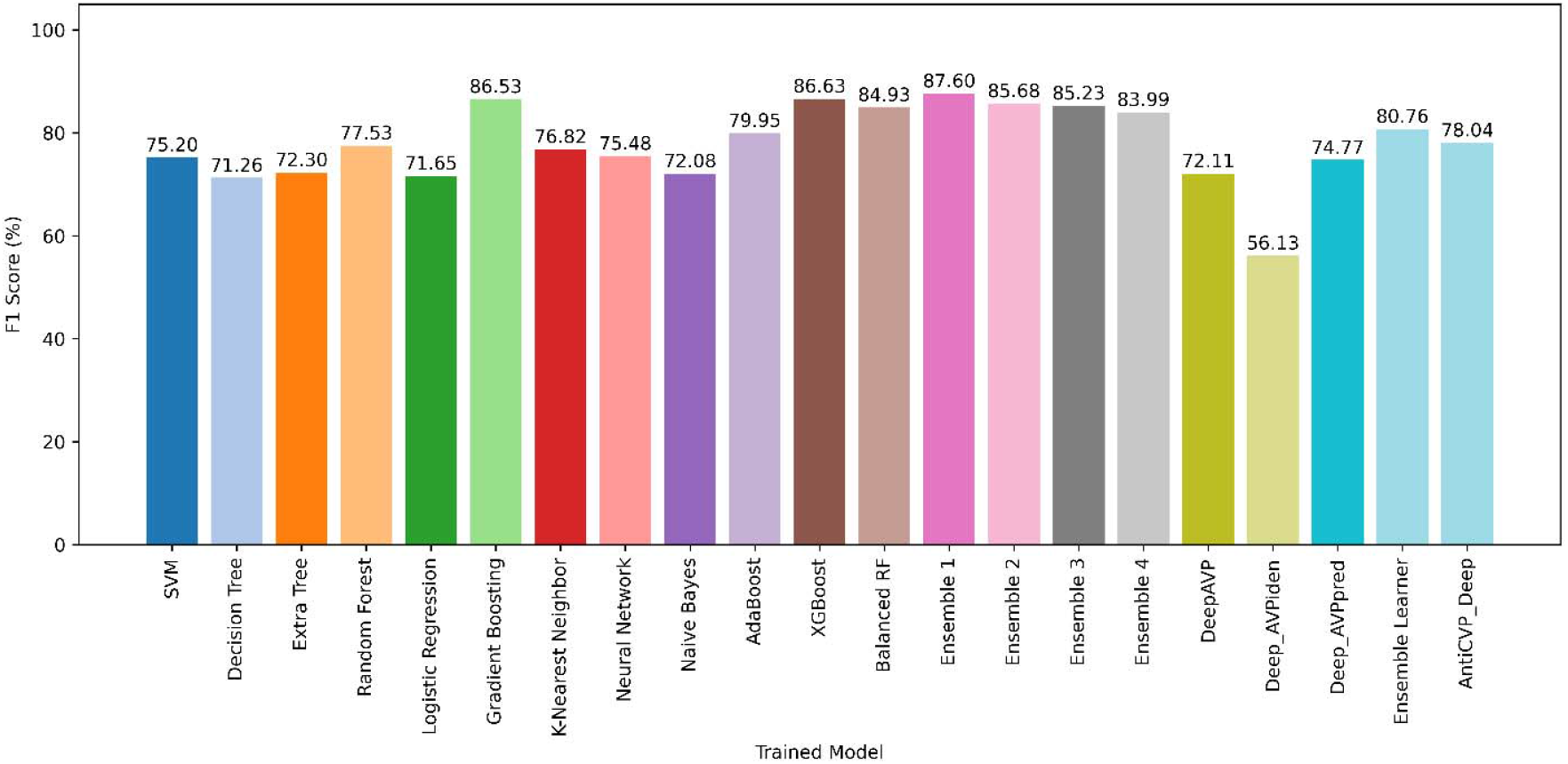
Comparison of the F1 scores. From the 21 models, the comparison depicts the highest F1 score found from Ensemble 1 model (dark pink line −87.6%). XGBoost and Gradient Boosting score the best from traditional machine learning models. They are shown by dark brown and light green lines with F1 scores of 86.63% and 86.53%, respectively. And the Deep learning models are shown to have underperformed in this comparison.

**Figure 7.**
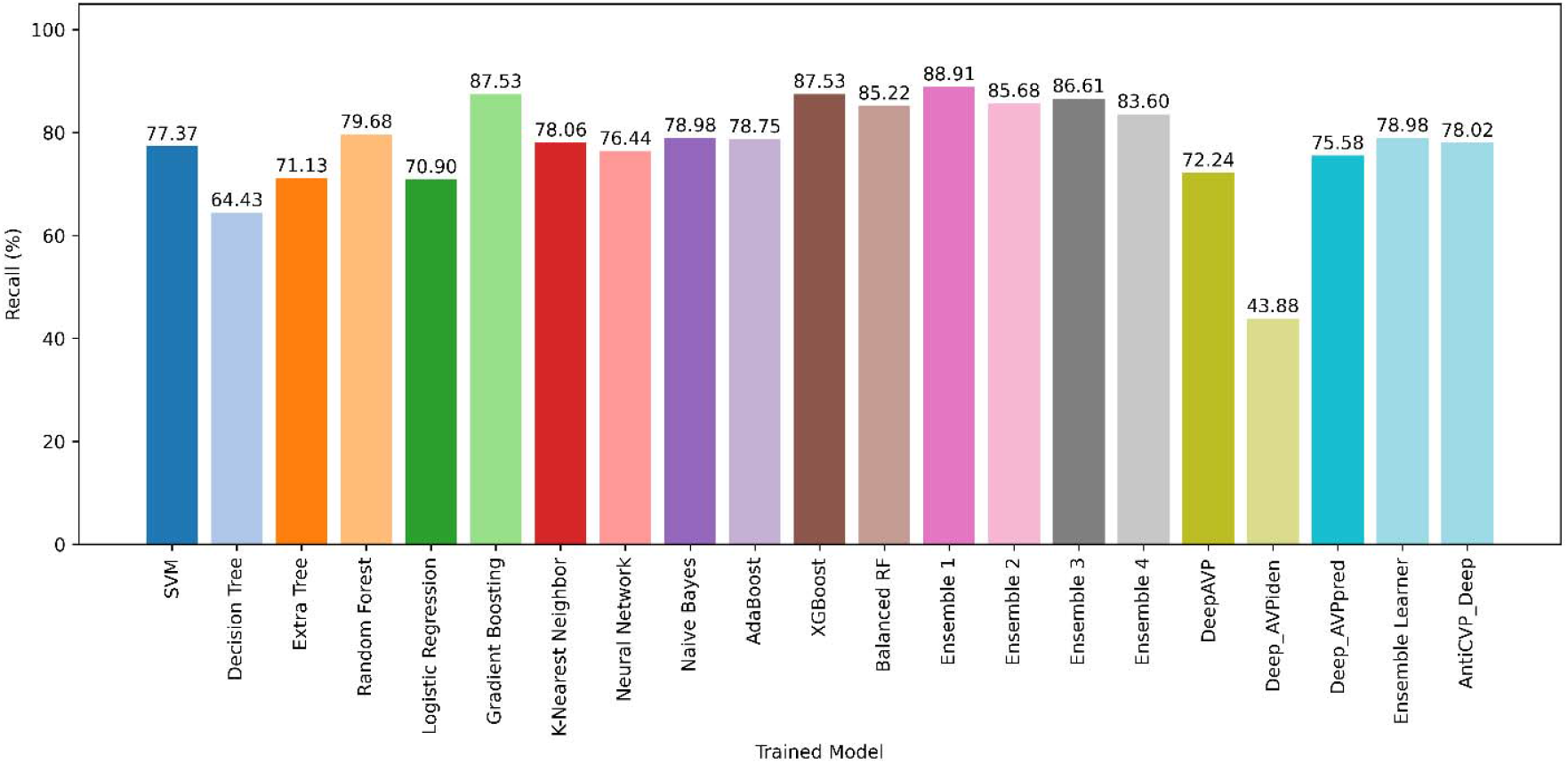
Comparison of the recall. The performance of each model in terms of recall is demonstrated here. The Ensemble 1 model scores the highest among all, with a recall score of 88.91%, shown in a dark pink line. From the traditional machine learning models, Gradient Boosting and XGBoost performed well; they both had an 87.53% recall value (light green and dark brown line, respectively).

**Figure 8.**
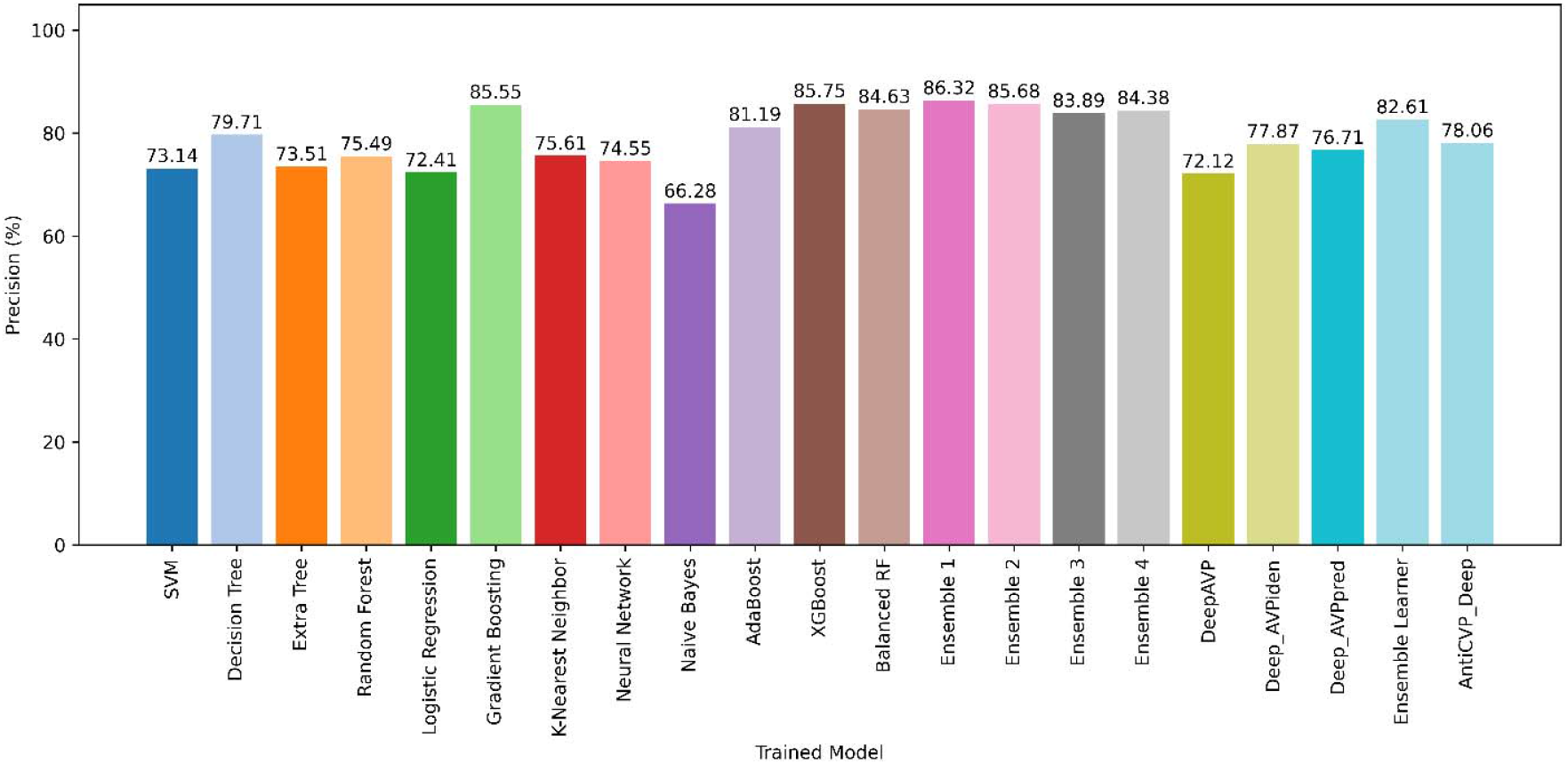
Comparison of the precision. The precision performance of each model is demonstrated in this figure. With a precision of 86.32%, the Ensemble 1 model shows the best score (dark pink line). The Ensemble 2 model also achieves the third-best precision output (85.68% - light pink line), showing the effectiveness of the constructed ensemble models.

In terms of the AUC metric, the Ensemble Learner achieved the highest score of 88.30%, while our proposed model closely follows with an AUC of 85.62% (**Figure 9**). Additionally, our proposed model recorded the lowest MAE at 14.01% (**Figure 11**) and the lowest RMSE of 37.43% (**Figure 12**). Our proposed model outperformed others in most metrics, showcasing their robustness and reliability.

**Figure 9.**
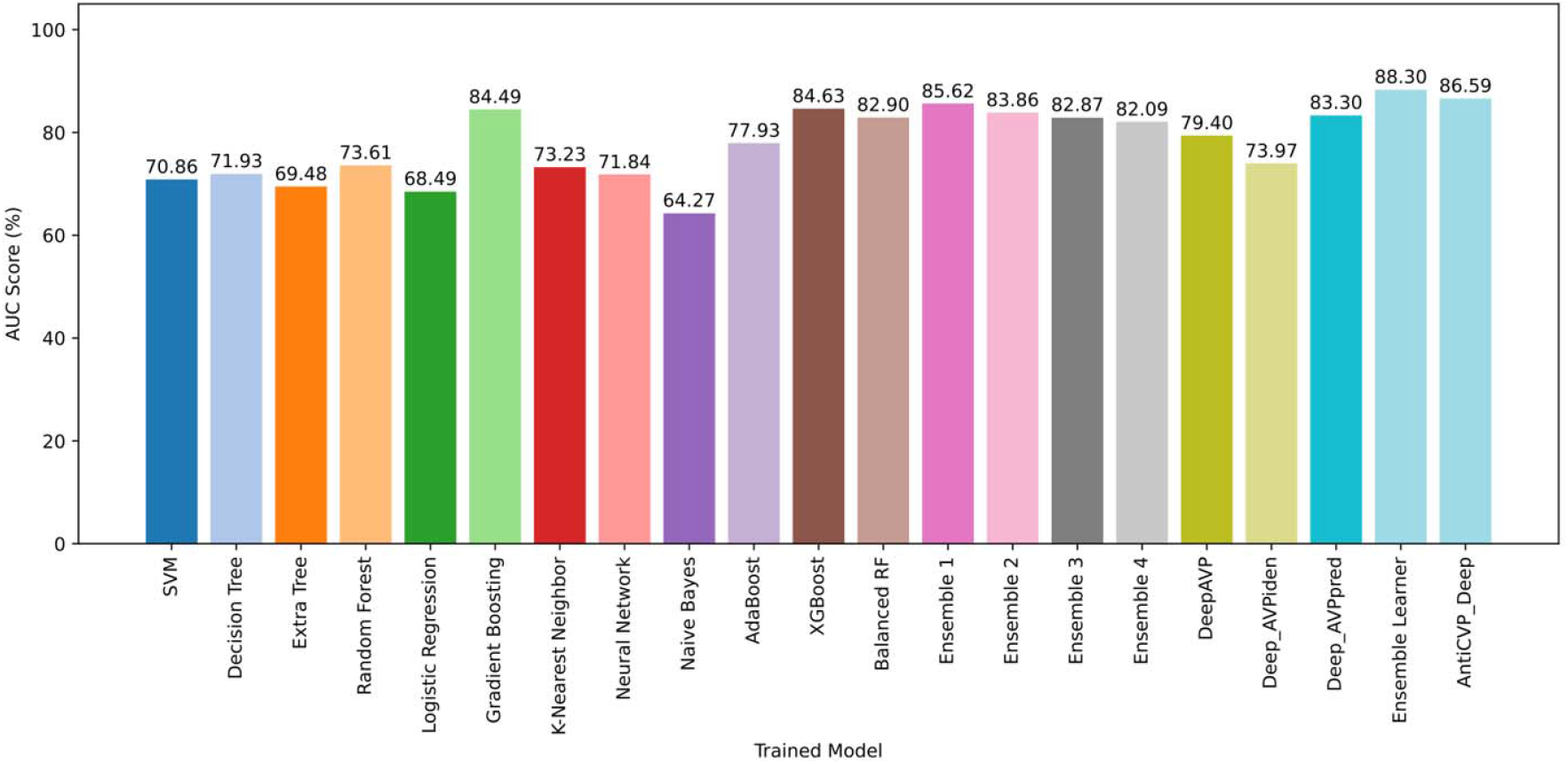
Comparison of the Area under ROC curve (AUC). The performance of each model in terms of AUC is demonstrated here. The Ensemble Learner model scores the highest among all, with an AUC of 88.3%, shown in a bright blue line. Among the other top performers, AntiCVP Deep and Balanced Random Forest achieved scores of 86.59% and 85.90%, represented by teal and orange lines, respectively. The proposed Ensemble 1 model ranks fourth with an AUC score of 85.62%, shown in deep purple.

**Figure 10.**
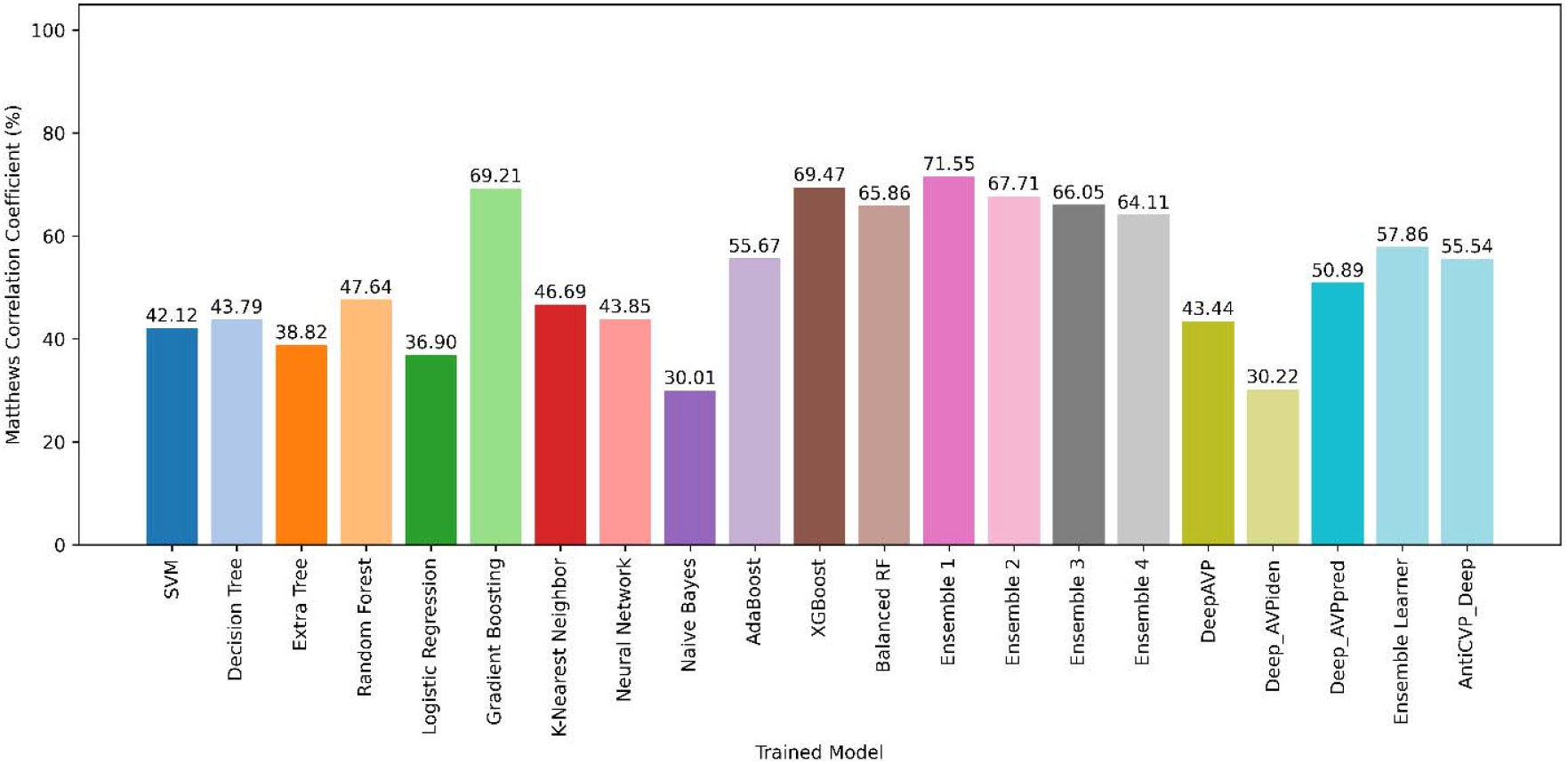
Comparison of the Matthews Correlation Coefficient. The MCC score of each model is demonstrated here. While the deep learning models like DeepAVP, Deep AVPiden lack to perform (43.44% - lime green and 30.22% - light lime green, respectively), the ensemble models achieve good scores. With an MCC score of 71.55%, the Ensemble 1 model is at the top in the overall comparison (dark pink line).

**Figure 11.**
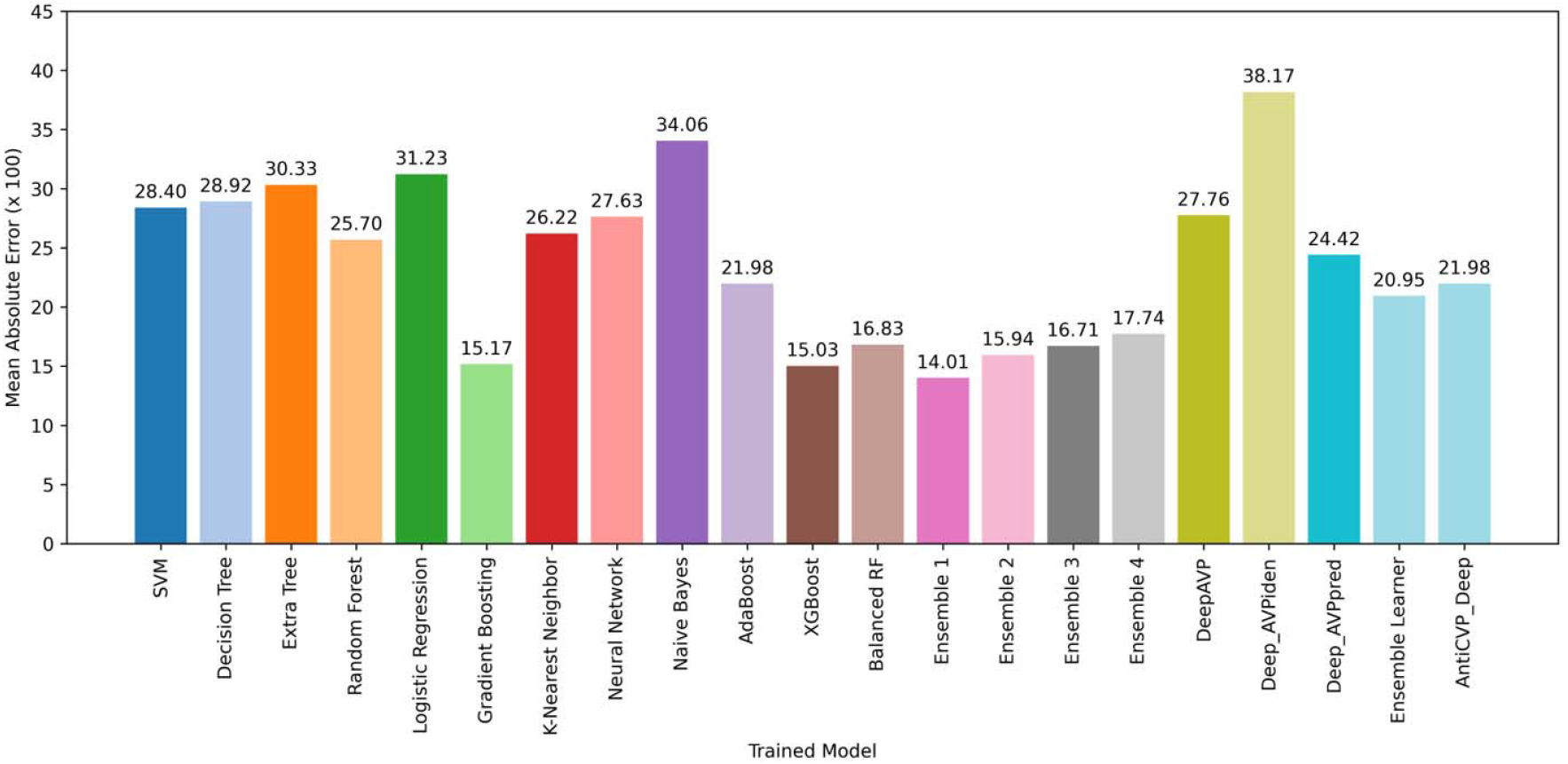
Comparison of the Mean Absolute Error. The MAE score of each model is demonstrated by different color lines. The Ensemble 1 model achieves the lowest MAE score (14.01 % - dark pink line), showing its effectiveness in predicting outcomes. XGBoost, Gradient Boosting are the winners among the traditional machine learning models (15.03% - dark brown and 15.17% - light green line, respectively). Subsequently, Ensemble 2 and 3 models show lower MAE scores (15.94% - light pink and 16.71% - dark gray line, respectively).

**Figure 12.**
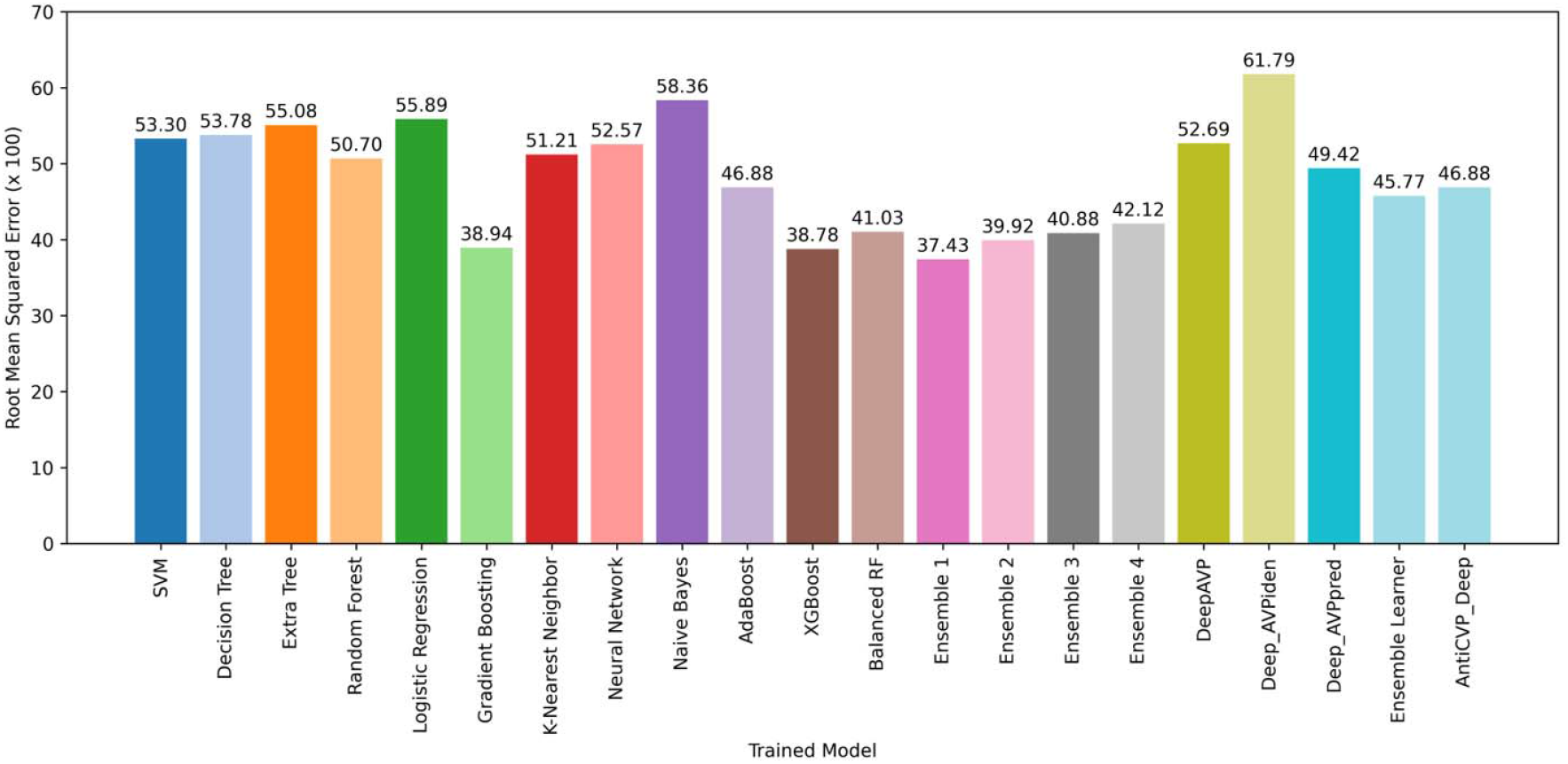
Comparison of the Root Mean Squared Error. The RMSE score of each model is demonstrated by different color lines. While the deep learning models like DeepAVP, Deep AVPiden achieve higher RMSE scores (52.96% - dark lime green and 61.79% - light lime green line, respectively), ensemble models gain the lower numbers. The ensemble 1 model attains 37.43%, which is the lowest among all models, as shown by the dark pink line here.

Additionally, **Figure 13** presents the Receiver Operating Characteristic (ROC) curves for all trained models evaluated on the test dataset. Among them our proposed model, consistently shows a higher true positive rate across different thresholds and achieves an overall superior ROC curve compared to the other methods, highlighting its robustness and effectiveness in capturing complex patterns in the data for accurate antiviral peptide prediction.

**Figure 13.**
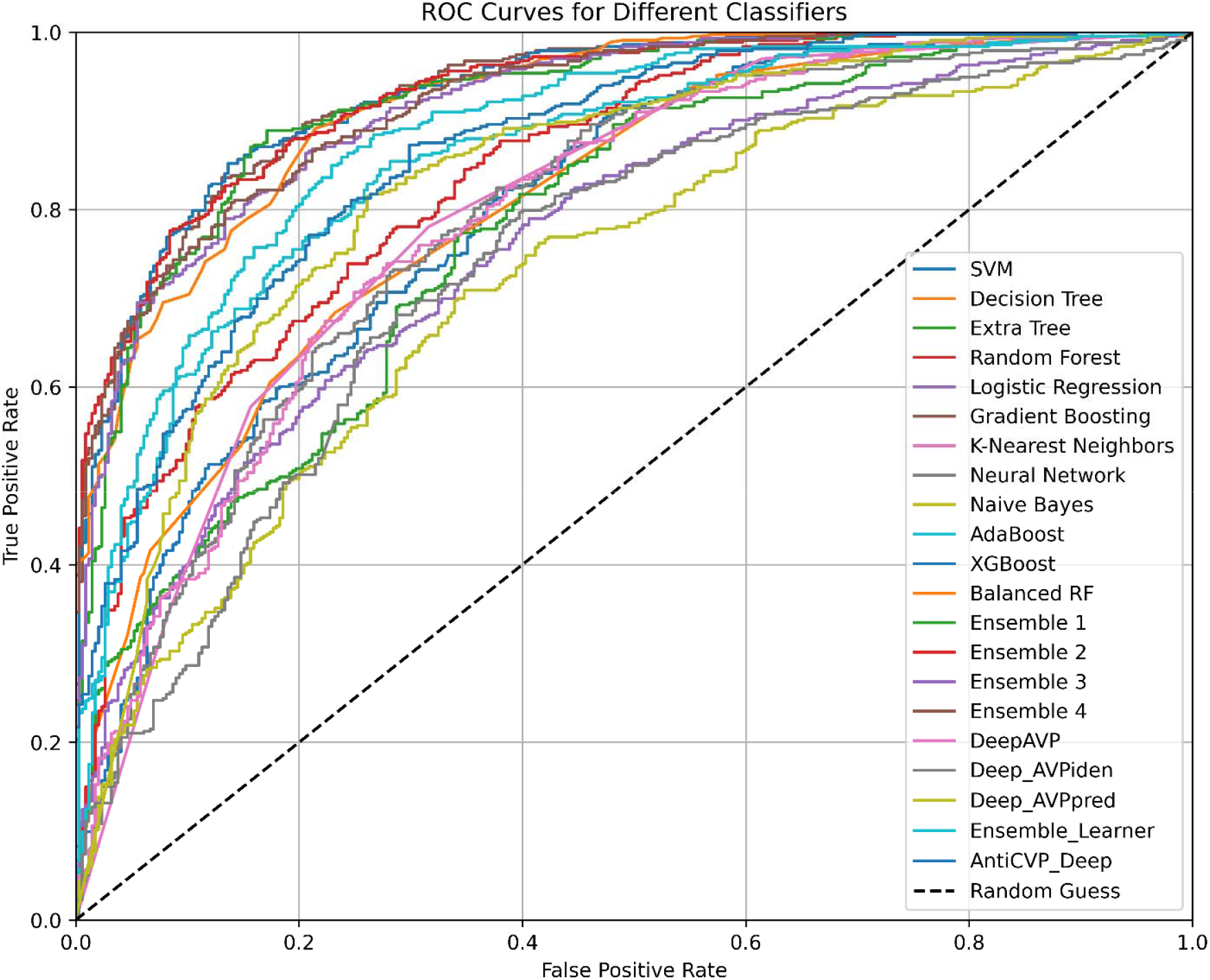
Comparison of Area Under the Curve (AUC) for all models. This is showing True Positive Rate (TPR) against the False Positive Rate (FPR).

The developed Ensemble 1 model was validated via 10-fold cross-validation, and metrics are expressed as mean ± standard deviation for each fold (**Figure 14**).

**Figure 14.**
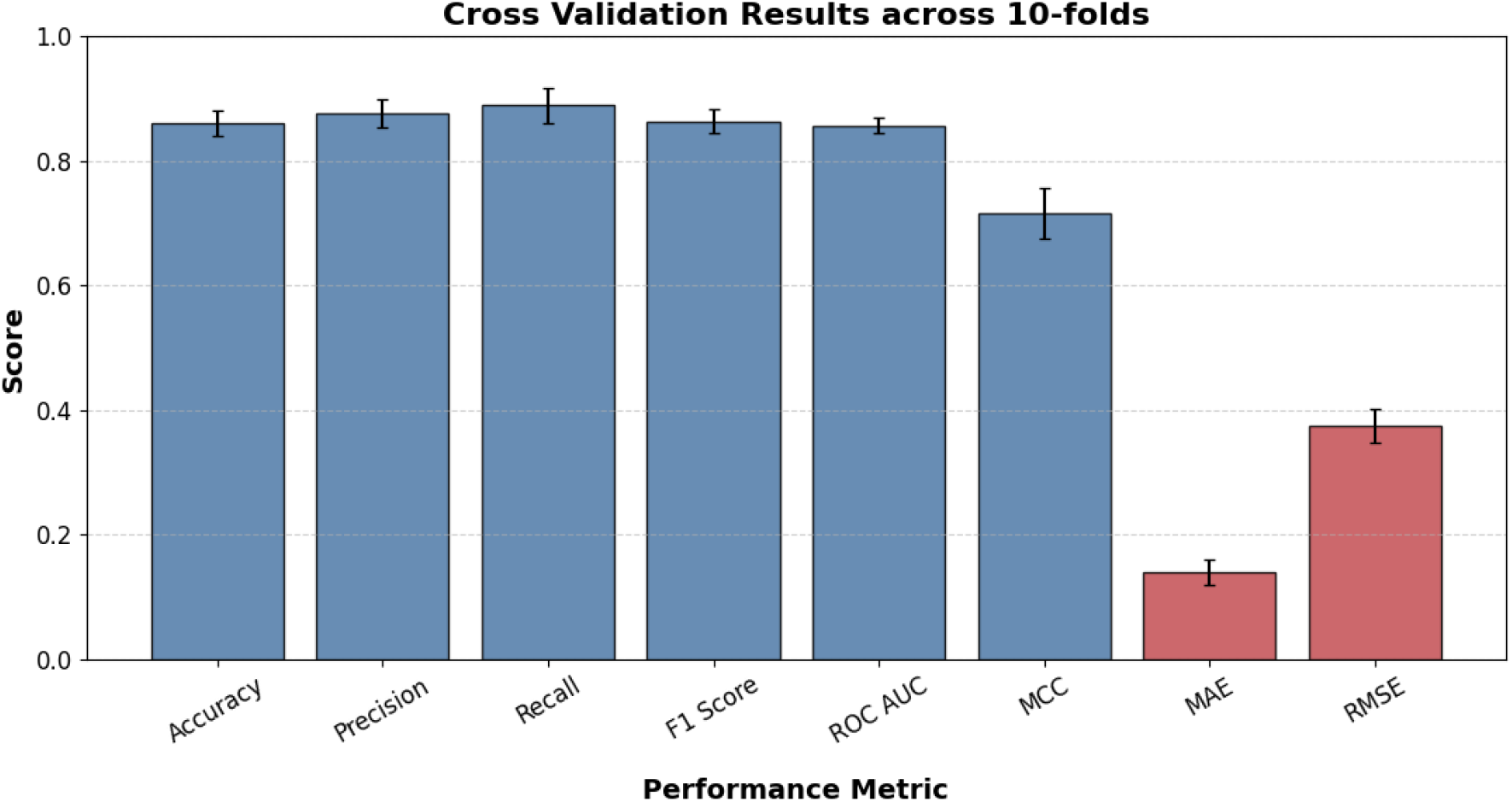
Performance of the Ensemble 1 model evaluated from 10-fold cross-validation with standard deviation. Classification metrics (blue): Accuracy (0.8599±0.0201), Precision (0.8760±0.0221), Recall (0.8891±0.0288), F1 Score (0.832±0.0192), ROC-AUC (0.8562±0.0125), MCC (0.7155±0.0407). Error metrics (red): MAE (0.1401±0.0201), RMSE (0.3743±0.0268).

## Discussion

Viral infections present a persistent threat to global health due to their lethality, exacerbated by the limits of conventional antiviral therapies, which include drug resistance, exorbitant costs, and extended development timelines. AVPs have significant advantages over traditional techniques due to their immunomodulatory qualities, ability to reduce resistance development, and capacity to enhance efficacy levels. Nonetheless, the identification of AVPs by experimental validation remains highly problematic due to the extensive time and costs involved, rendering computational predictions of AVPs essential for advancing the field.

The suggested model APDeeM has shown enhanced performance (**Supplementary Table 4**) in identifying AVPs due to its integration with multiple models (**Figure 1**), including Gradient Boosting, Random Forest, K-Nearest Neighbors (KNN), and AdaBoost. It attained a commendable accuracy of 85.99% and an F1 score of 87.6%, indicating its superiority over existing models (**Supplementary Table 4, Figure 5** and **Figure 6**). This ensures that the model is robust, accurate, and dependable in its ability to identify both non-antiviral and antiviral peptides, underscoring the importance of minimizing the impact of false positives and false negatives. In addition to emphasizing the model’s minimal misclassification rate, which is essential for the reliability of antiviral peptide detection, the model’s precision (86.32%) and recall (88.91%) further validate its accuracy in identifying true positives of antiviral peptides (**Supplementary Table 4, Figure 7** and **Figure 8**).

Additionally, true positive rates of other models were illustrated and their evaluation metrics were achieved (**Table 1** and **Supplementary Table 4**).

The proposed Ensemble Antiviral Peptide Detection model (APDeeM), which integrates Gradient Boosting, Random Forest, K-Nearest Neighbors (KNN), and AdaBoost, demonstrated improved efficacy in identifying antiviral peptides (AVPs). It achieved an accuracy of 85.99% and an F1 score of 87.6% (**Supplementary Table 4, Figure 5**, and **Figure 6**). Gradient Boosting incrementally enhances accuracy by correcting prior errors, Random Forest mitigates overfitting, KNN provides a proximity-based classification approach, and AdaBoost focuses on difficult instances. The model captures diverse data patterns through the voting mechanism, leading to stronger generalization and predictive performance. The performance of APDeeM surpasses that of the current model for antiviral peptide detection employed in this research.

Consequently, it yields precise results and can reduce the occurrence of false positives and false negatives in the identification of both antiviral and non-antiviral peptides. The APDeeM tools accurately identify true positive AVP, with a precision of 86.32% and a recall of 88.9%, with a little misclassification rate (**Supplementary Table 4, Figure 7** and **Figure 8**). The incorporation of gradient boosting into the APDeeM yields the highest precision (86.32%), ensuring the effective identification of real negatives and substantially improving the model’s capacity to minimize false positives.

This enhancement and performance emphasize its advantages over current models that depend on labor-intensive laboratory experiments and are subject to data imbalance and restricted feature sets. The APDeeM effectively addresses the significant gap in AVP identification by utilizing a comprehensive dataset comprising 14 distinct features and implementing advanced preprocessing techniques, including the Synthetic Minority Oversampling Technique (SMOTE), to rectify class imbalance, thereby offering a more efficient and scalable alternative to conventional methods.

Moreover, the APDeeM model attained a notable Matthews Correlation Coefficient (MCC) of 71.55%, highlighting its balanced efficacy in differentiating between antiviral and non-antiviral peptide identification (**Supplementary Table 4** and **Figure 11**). This update provides a substantial enhancement over the current models, as the previous models encounter difficulties with imbalanced datasets. The low Mean Absolute Error (MAE) of 14.01% and Root Mean Squared Error (RMSE) of 37.43% further substantiate the dependability and accuracy of the APDeeM model for AVP identification (**Supplementary Table 4, Figure 12** and **Figure 13**). To address the limitations of previous studies on AVP detection, which frequently encountered restricted features and data imbalance, the APDeeM utilized a diverse array of unique peptide characteristics and incorporated an ensemble method to learn complex patterns, representing a significant advancement. Also, the model performs consistently, as seen by the low standard deviations for all assessment criteria **(Figure 14)**. This shows that during each fold of the cross-validation, the model is tested on randomly chosen subsets and remains stable and credible.

The improved accuracy of APDeeM compared to prior models employed on the current dataset is mostly due to the incorporation of 14 distinct peptide characteristics that offer a thorough biochemical characterization of the AVPs. The characteristics such as molecular weight, hydrophobicity (GRAVY), net charge, and the Boman index are essential for the interactions between peptides and viral components.

For example, highly charged peptides form strong contact with the membranes, while hydrophobic residues affect membrane penetration and stability. These characteristics within the machine learning framework indicate that the model encapsulates fundamental biological factors linked to antiviral action. However, introduction of intricate features may hinder the model’s utility in more advanced or beginner-level settings despite the importance. Future improvements should be focused on enhancing the model to have a balanced performance so that it overcomes the limitations and potentially useful to researchers and clinicians through web-based or application-based platforms.

## Conclusion

The present study describes the development of the APDeeM framework and evaluates its performance across eight comprehensive metrics. In addition, the proposed model is compared with other existing approaches to highlight its relative advantages. We conclude by outlining the potential applications of the framework in antiviral research and propose future perspectives for its further development and enhancement. We anticipate that this study will significantly impact the discovery of antiviral peptides (AVPs), offering a new computational avenue for the development of highly efficient, accurate, and reliable methods for identifying unknown AVPs. Consequently, this approach may accelerate the formulation of antiviral therapies—an essential factor in emergency response during viral outbreaks. Moreover, the integration of machine learning techniques with 14 distinct peptide characteristics ensures the development of a powerful and robust model capable of offering valuable insights into the complex attributes of AVPs. Overall, the findings of the present work contribute to the advancement of computational strategies in the fight against viral infections, ultimately supporting global public health initiatives.

## Ethics Statement

This study did not involve human participants, animal subjects, or any clinical trials. Therefore, ethical approval was not required. No personal or sensitive information was collected or used for this research.

## Data Availability Statement

The data underlying this article will be shared on reasonable request to the corresponding author.

## Author contributions statement

**Mohammad Uzzal Hossain:** Data curation, Formal analysis, Conceptualization, Methodology, Visualization, Writing – original draft. **Md. Romzan Alam:** Data curation, Formal analysis, Conceptualization, Methodology, Visualization, Writing – original draft. **SM Sajid Hasan:** Data curation, Formal analysis, Conceptualization, Methodology, Visualization, Writing – original draft. **Mohammad Nazmus Sakib:** Data curation, Methodology, Visualization, Writing – original draft. **Marjia Akter Suchi:** Data curation, Formal analysis, Visualization. **Zeba Sanjida:** Data curation, Formal analysis, Visualization. **A.B.Z. Naimur Rahman:** Data curation, Visualization, Formal analysis. **Arittra Bhattacharjee:** Data curation, Validation, Visualization, Writing – review & editing. **Zeshan Mahmud Chowdhury:** Data curation, Validation, Writing – review & editing. **Ishtiaque Ahammad:** Data curation, Validation, Writing – review & editing. **Muhammad Aminur Rahman:** Conceptualization, Investigation, Resources, Supervision, Writing – review & editing, Data curation, Formal analysis. **Saiful Azad:** Conceptualization, Investigation, Resources, Supervision, Writing – review & editing, Data curation, Formal analysis. **Md. Salimullah:** Conceptualization, Investigation, Resources, Supervision, Writing – review & editing, Data curation, Formal analysis.

## Acknowledgments

This research is the result of a collaborative effort between Green University of Bangladesh and the National Institute of Biotechnology, Bangladesh. Both institutions have contributed equally to the conception, development, and execution of the work. The authors gratefully acknowledge the support and cooperation provided by both organizations throughout the course of this study.

