## Supplementary Figures for "APDeeM: A machine Learning strategy towards Effective Peptide Vaccine Candidates Identification against Different Types of Viruses"

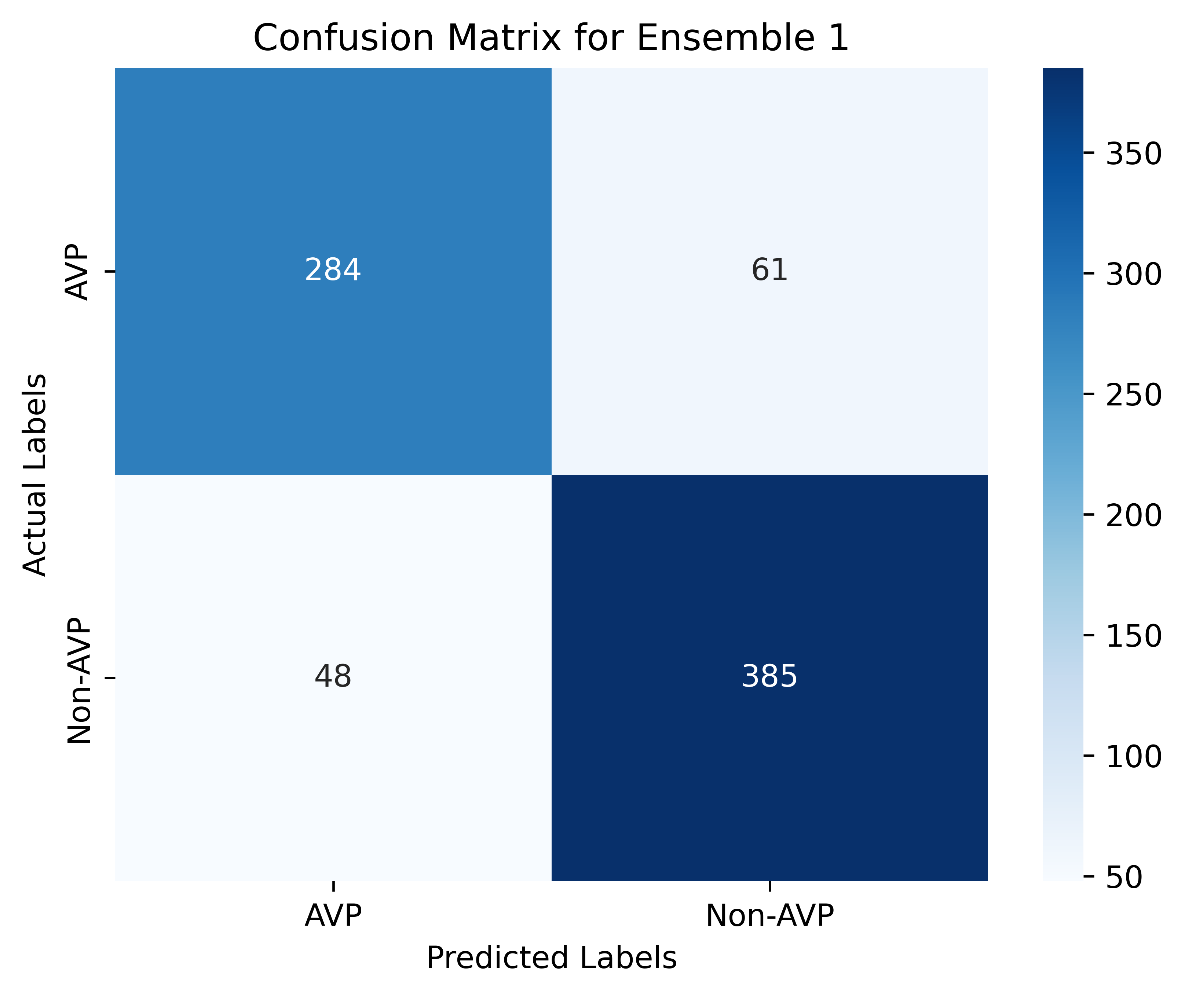

**Supplementary Figure 1: Confusion matrix of the Ensemble 1 model.**

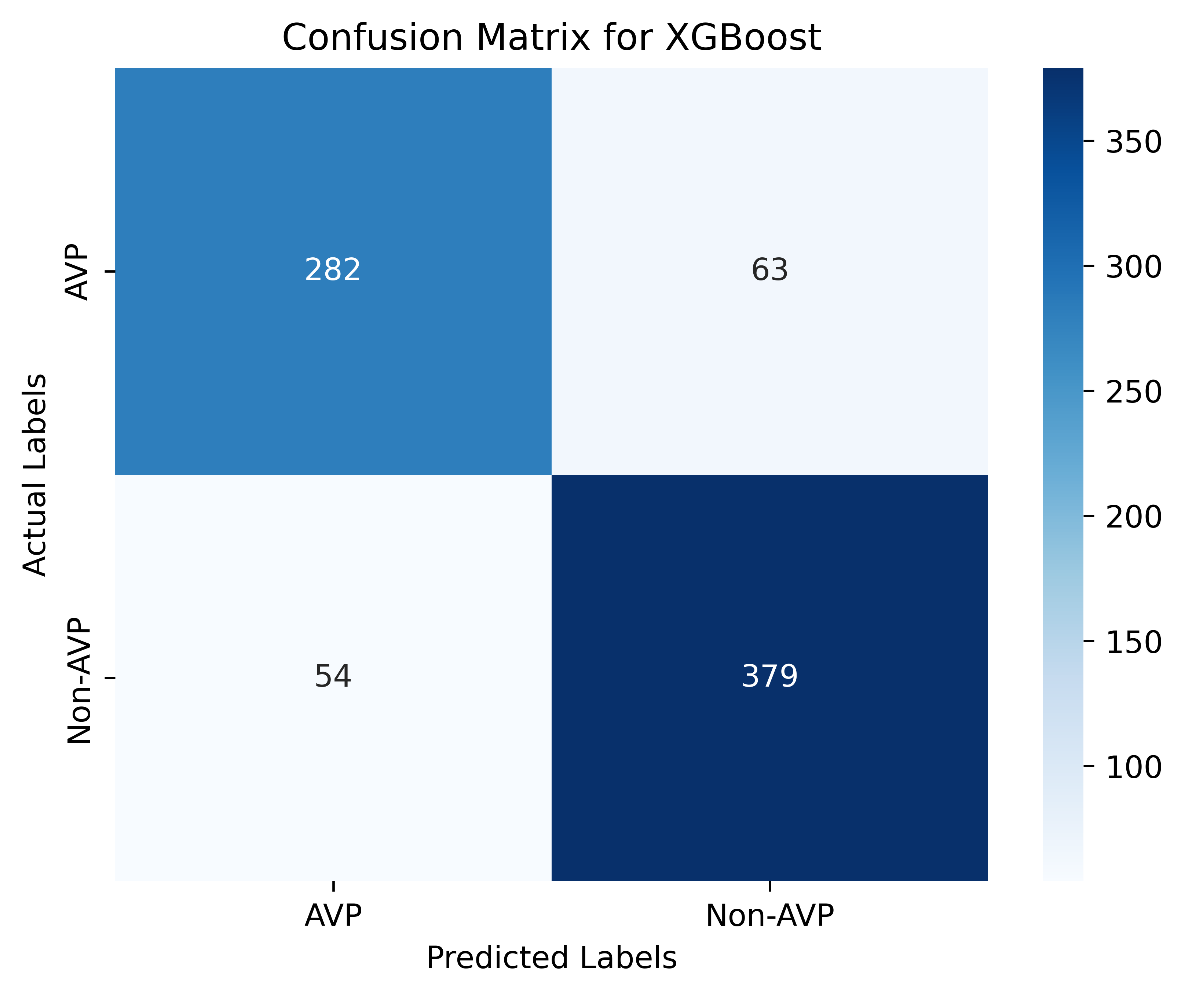

**Supplementary Figure 2: Confusion matrix of the XGBoost model.**

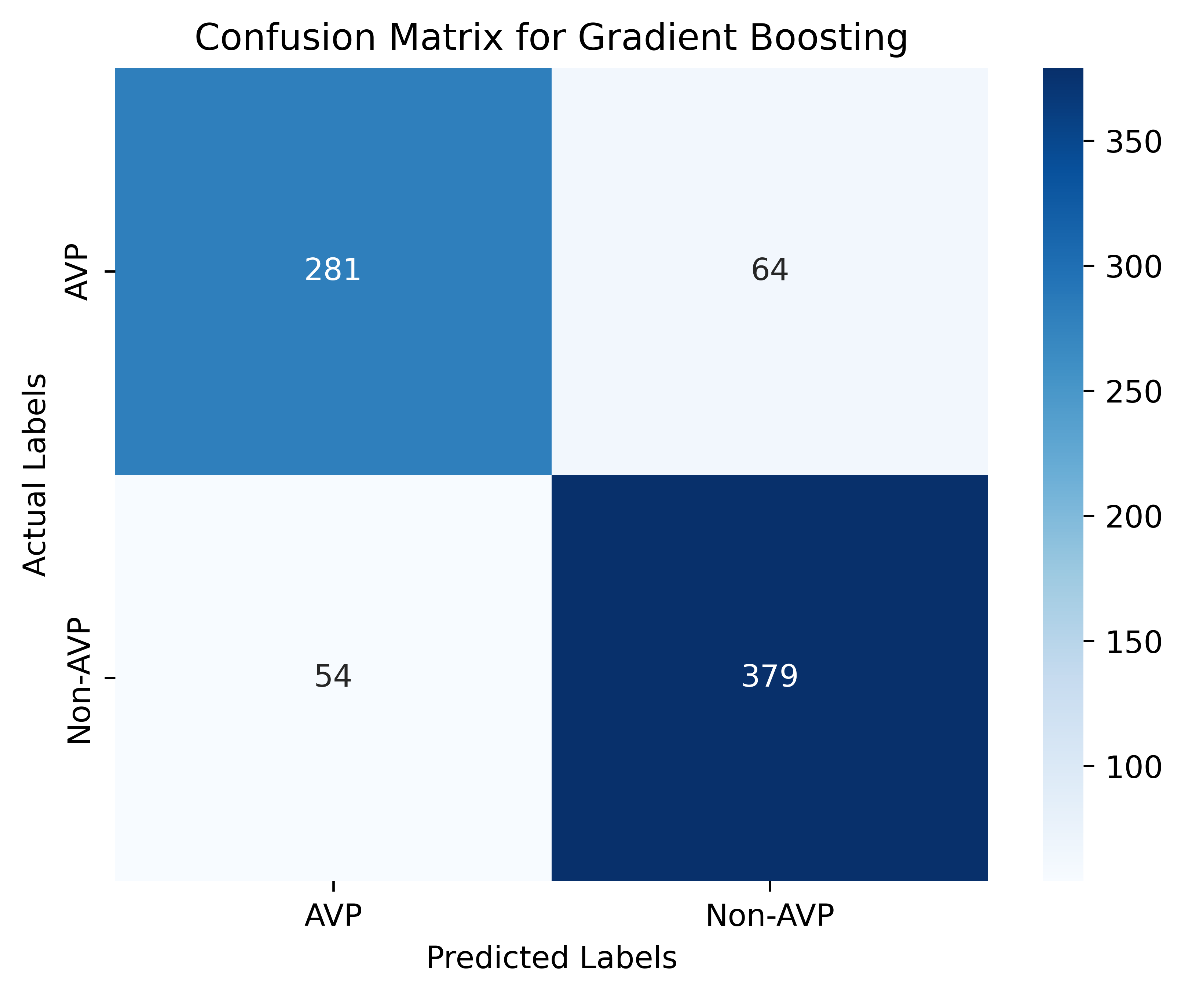

**Supplementary Figure 3: Confusion matrix of the Gradient Boosting model.**

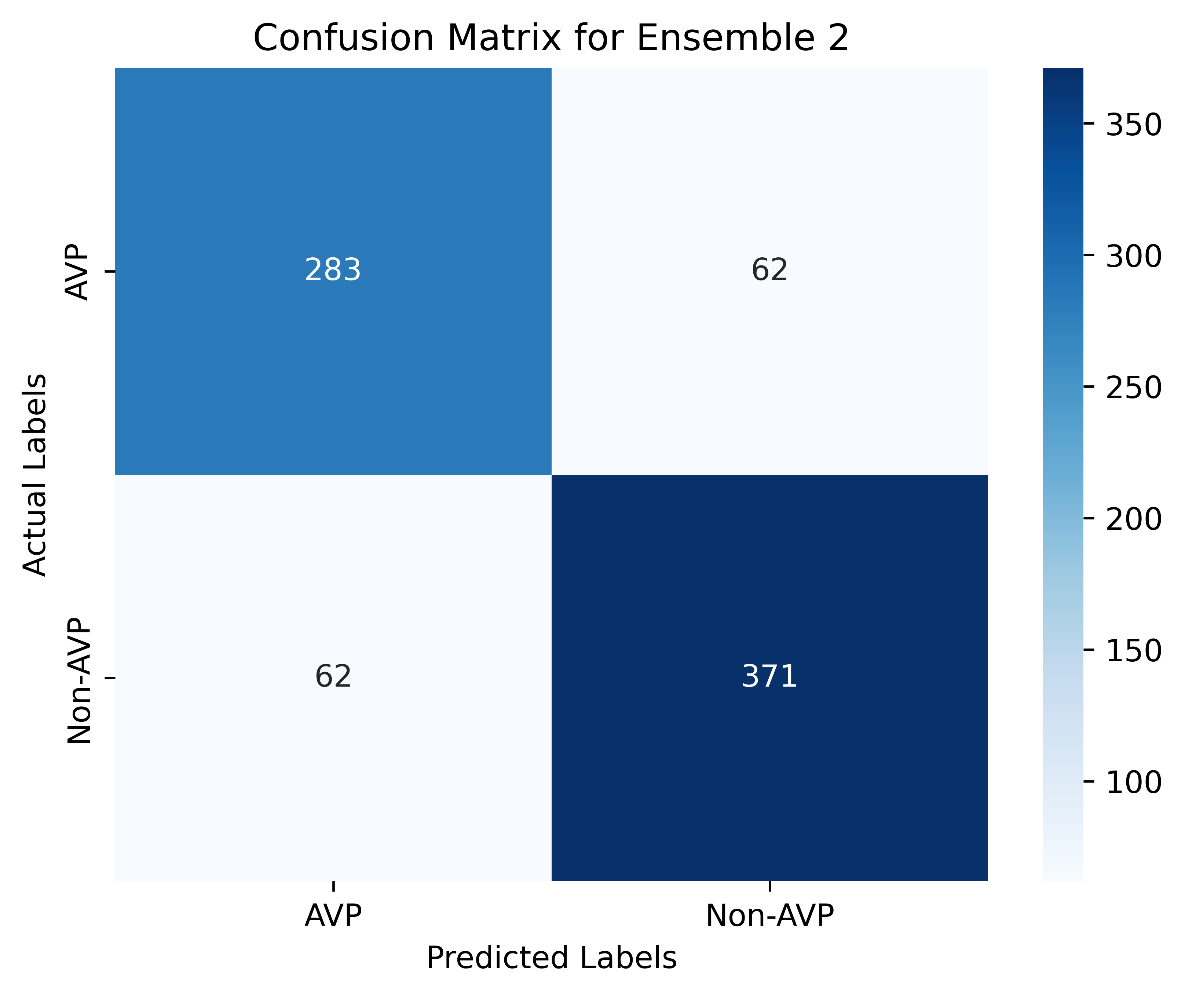

**Supplementary Figure 4: Confusion matrix of the Ensemble 2 model.**

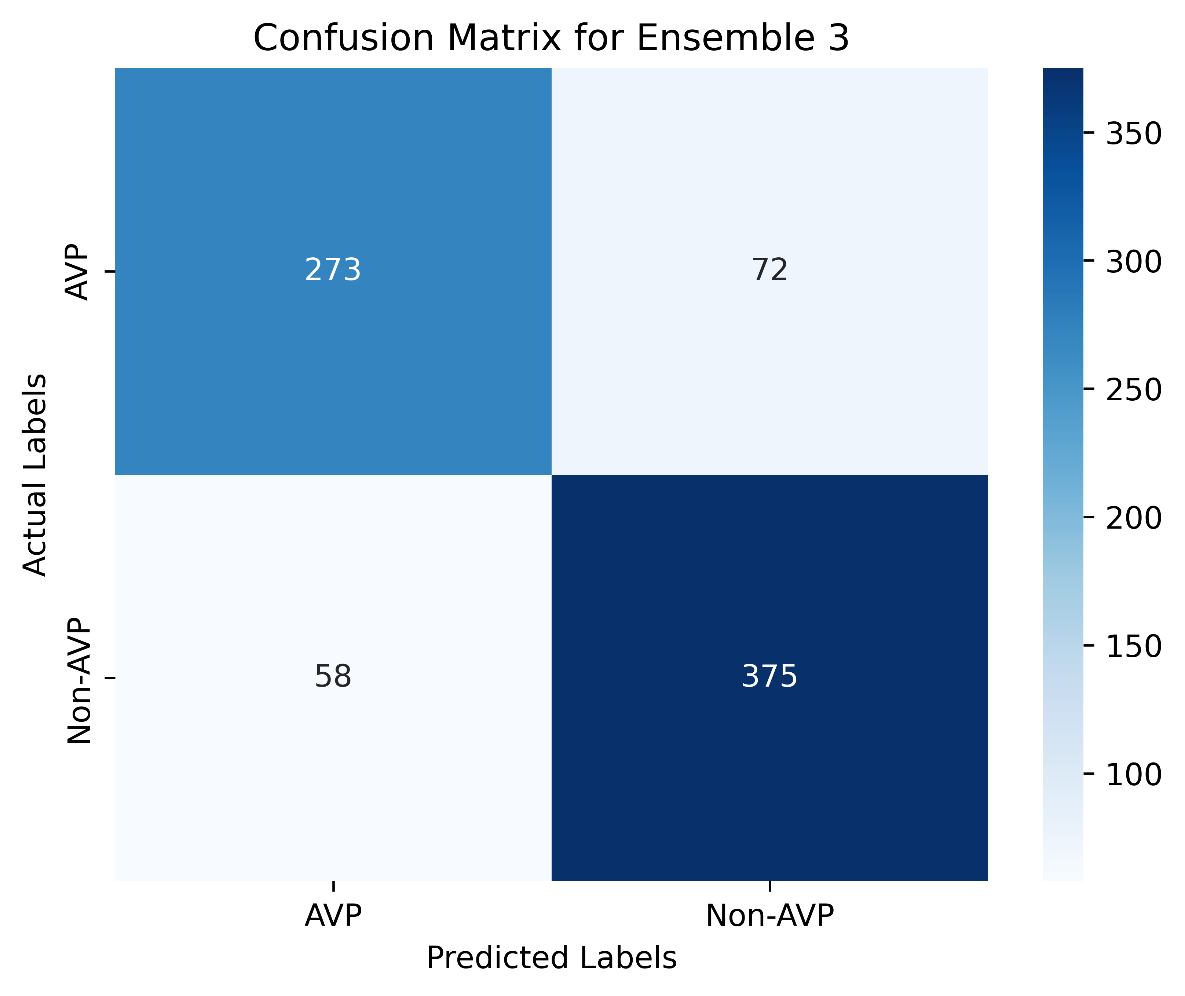

**Supplementary Figure 5: Confusion matrix of the Ensemble 3 model.**

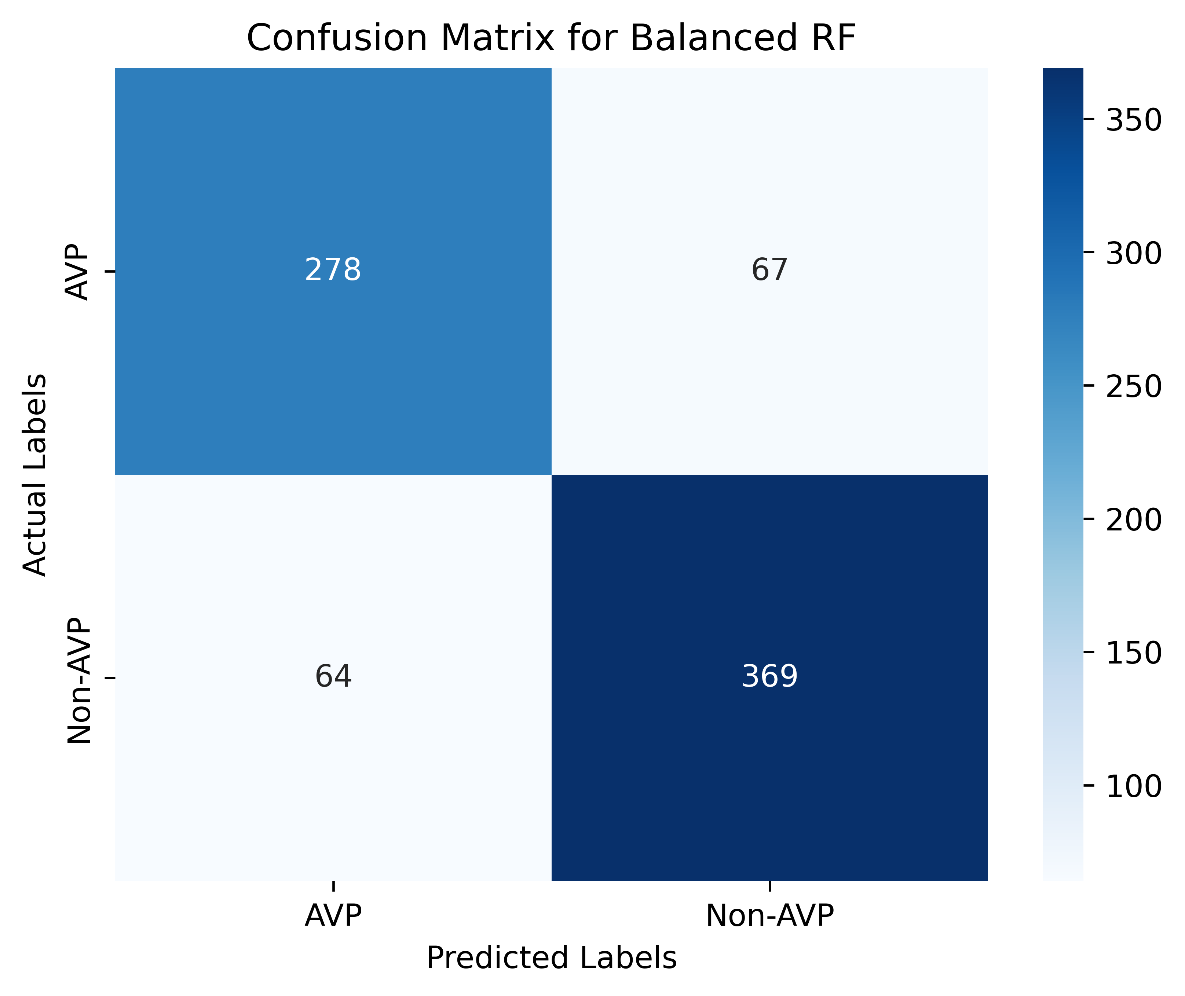

**Supplementary Figure 6: Confusion matrix of the Balanced Random Forest model.**

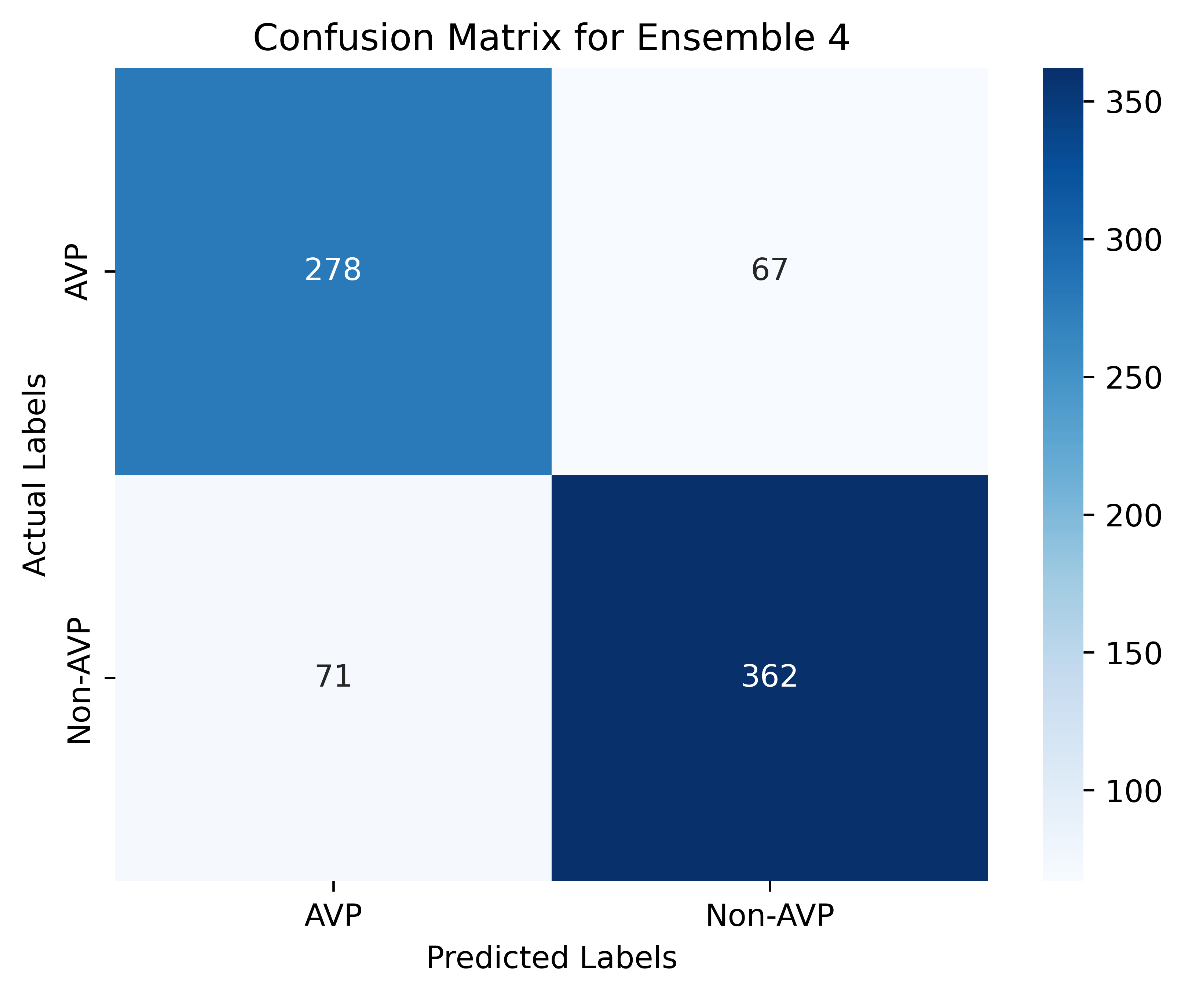

**Supplementary Figure 7: Confusion matrix of the Ensemble 4 model.**

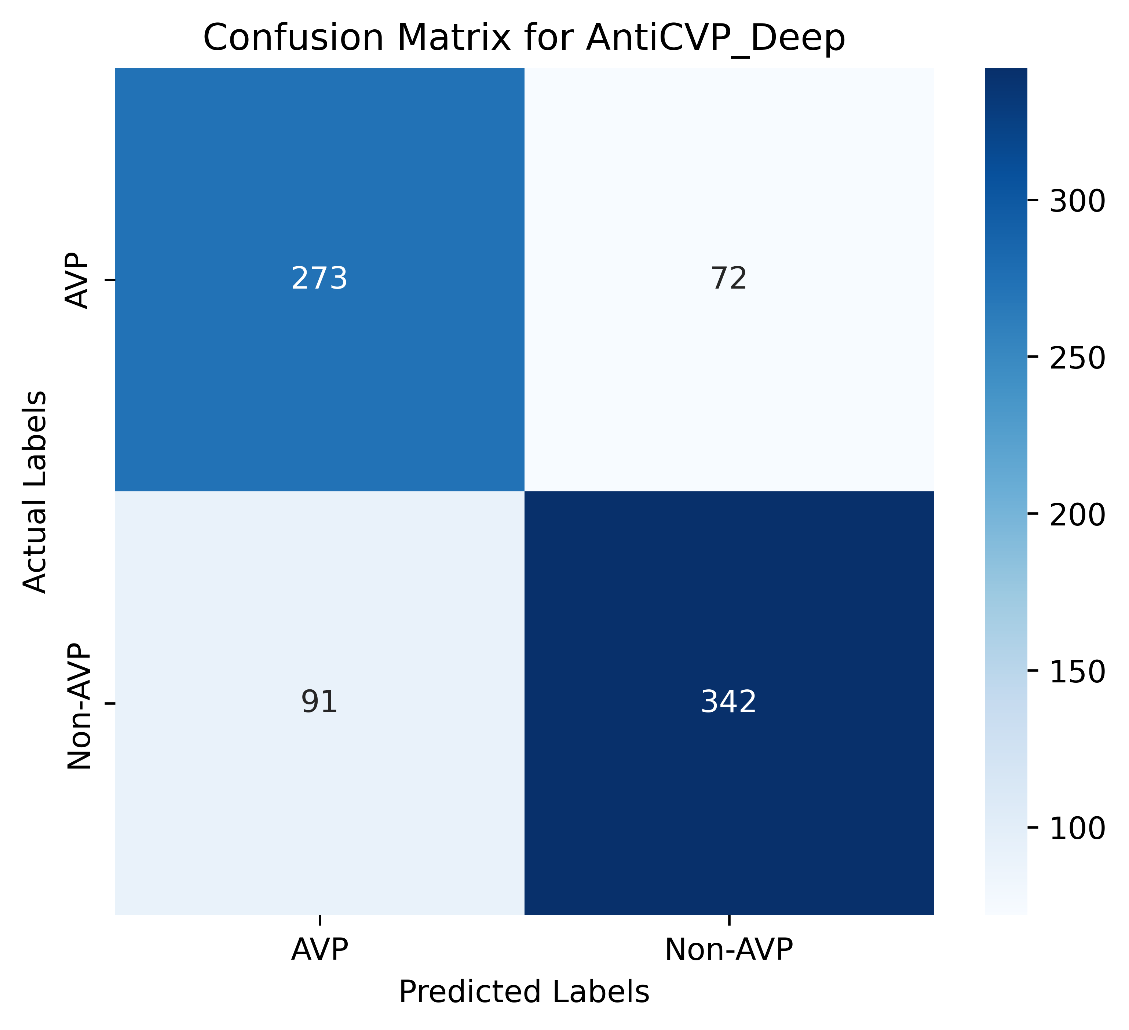

**Supplementary Figure 8: Confusion matrix of the Ensemble Learner model.**

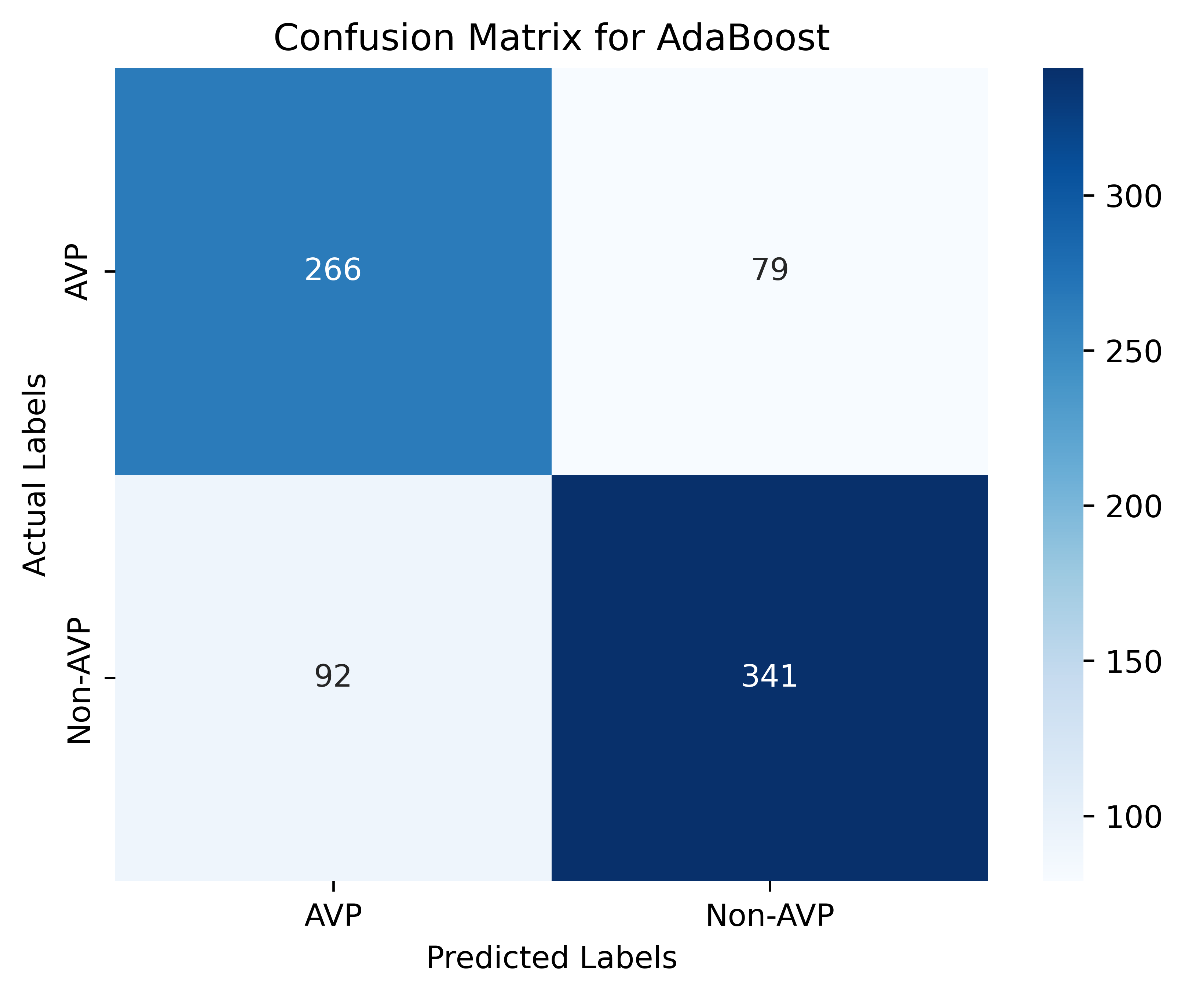

**Supplementary Figure 9: Confusion matrix of the AdaBoost model.**

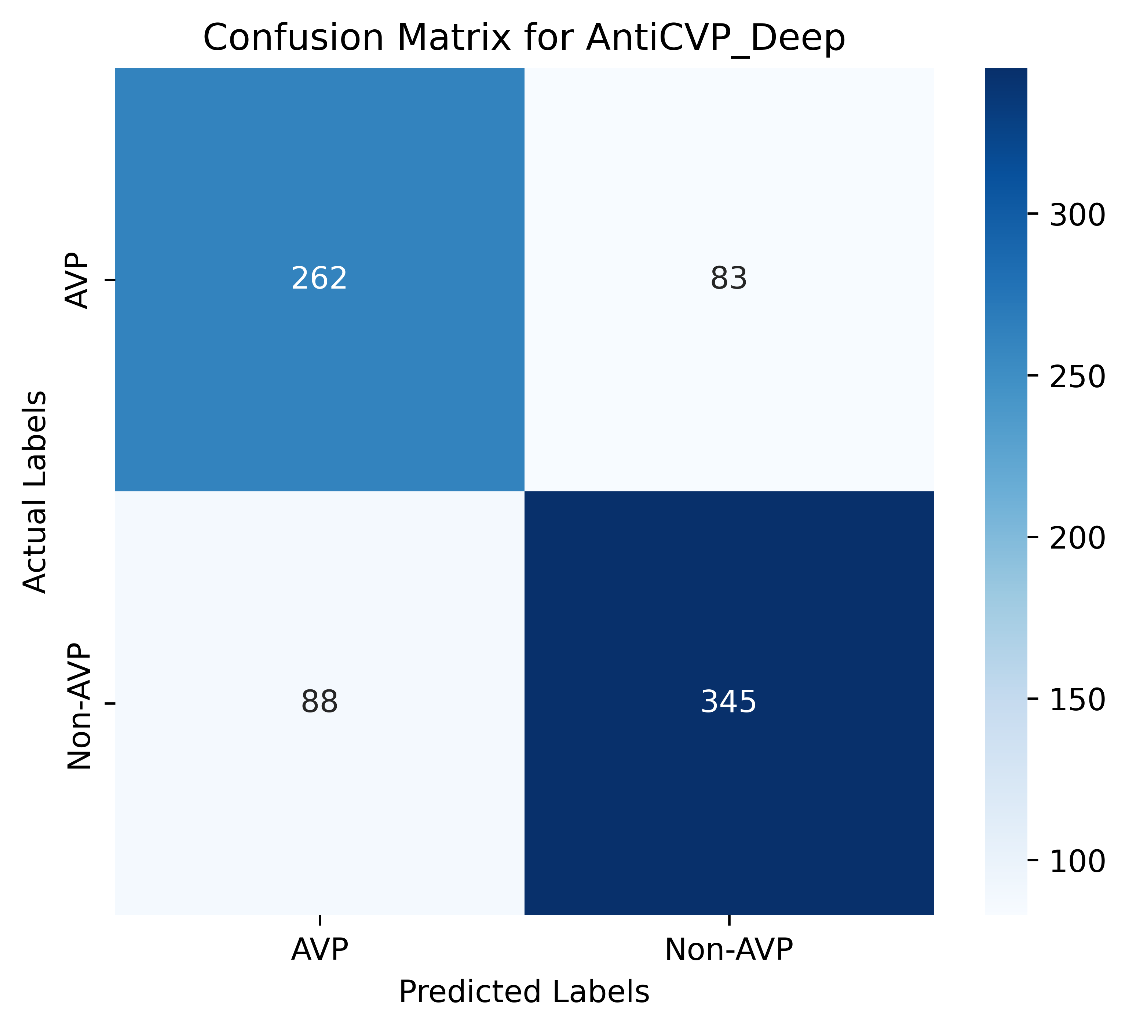

**Supplementary Figure 10: Confusion matrix of the AntiCVP_Deep model.**

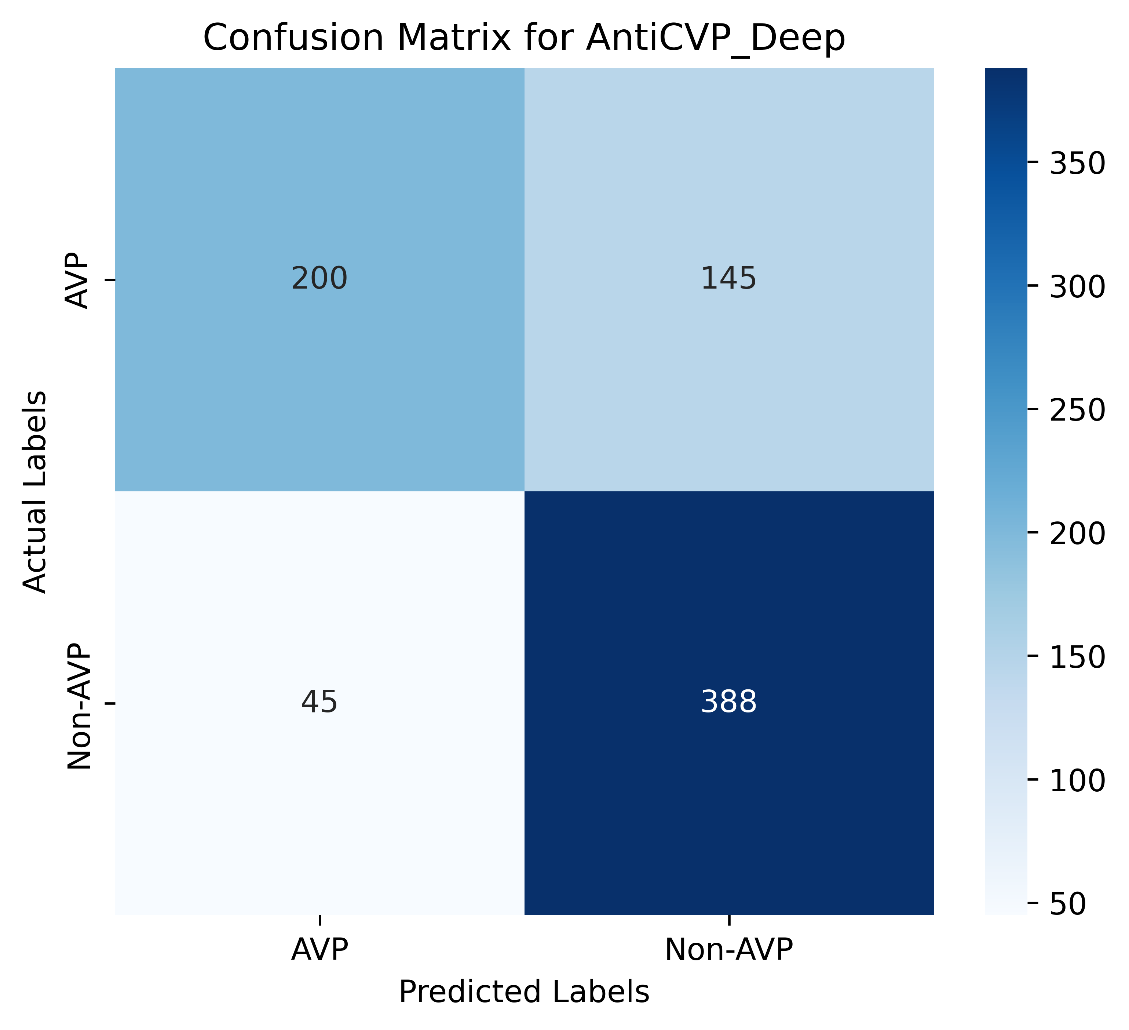

**Supplementary Figure 11: Confusion matrix of the Deep_AVPpred model.**

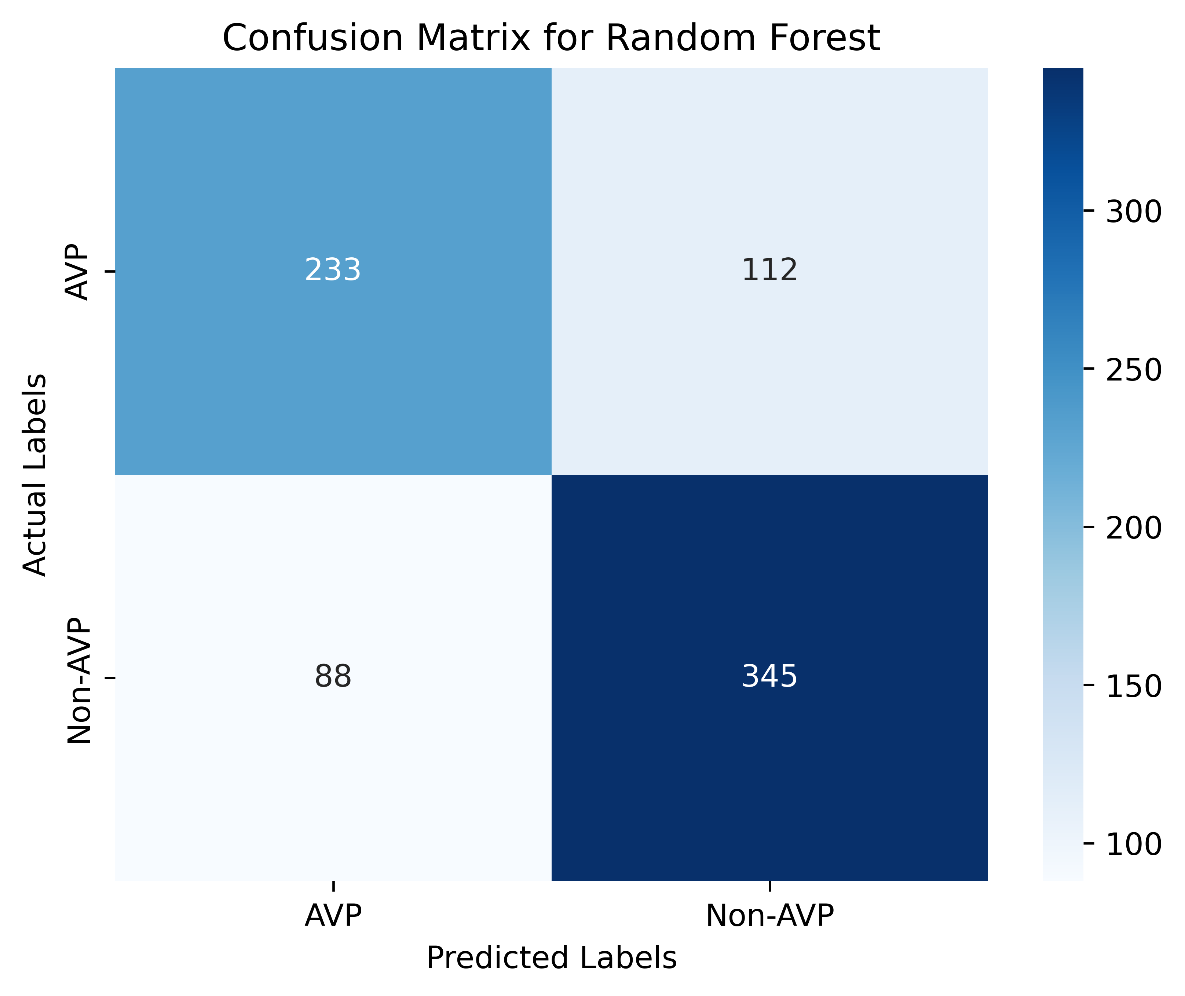

**Supplementary Figure 12: Confusion matrix of the Random Forest model.**

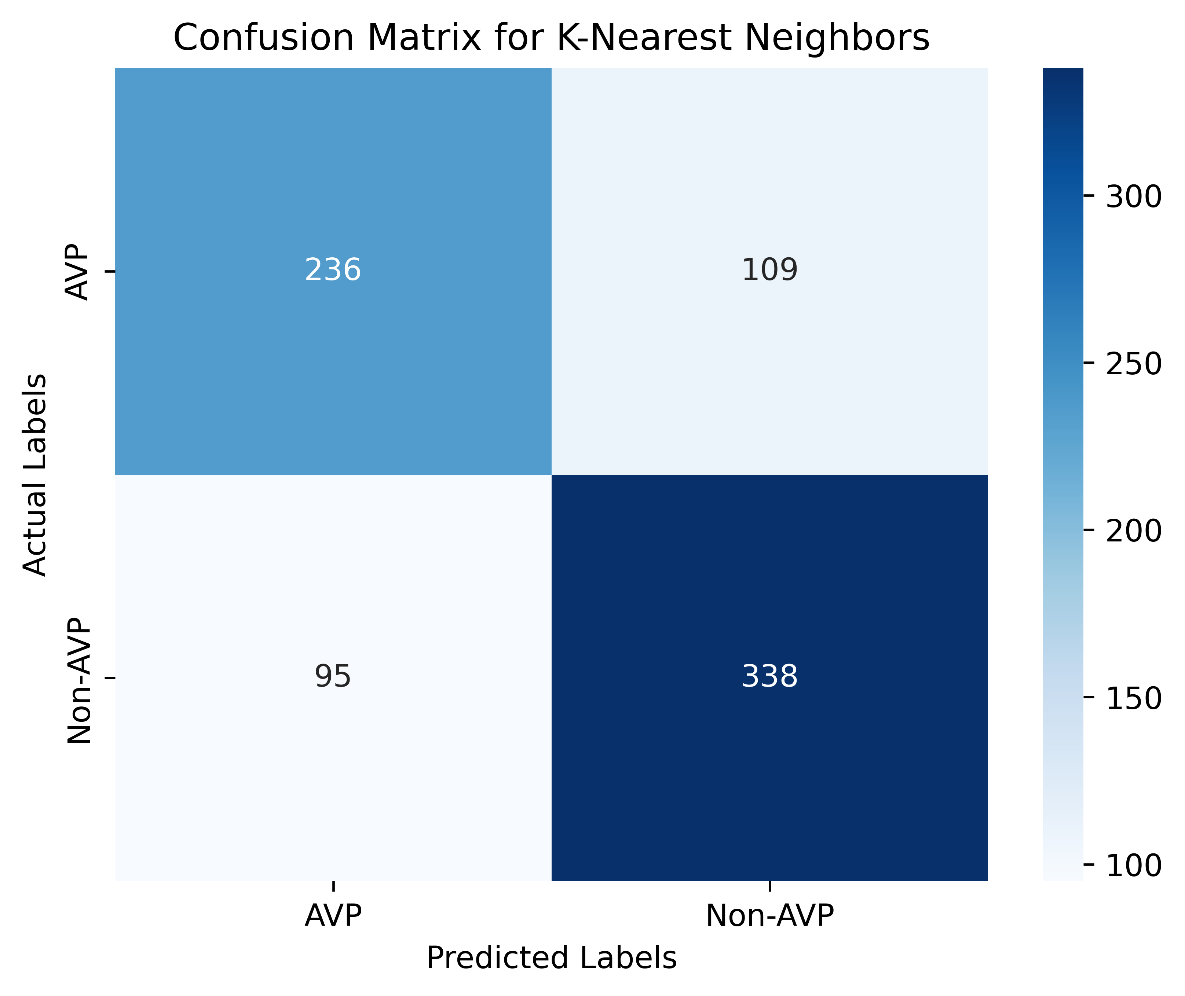

**Supplementary Figure 13: Confusion matrix of the K-Nearest Neighbors model.**

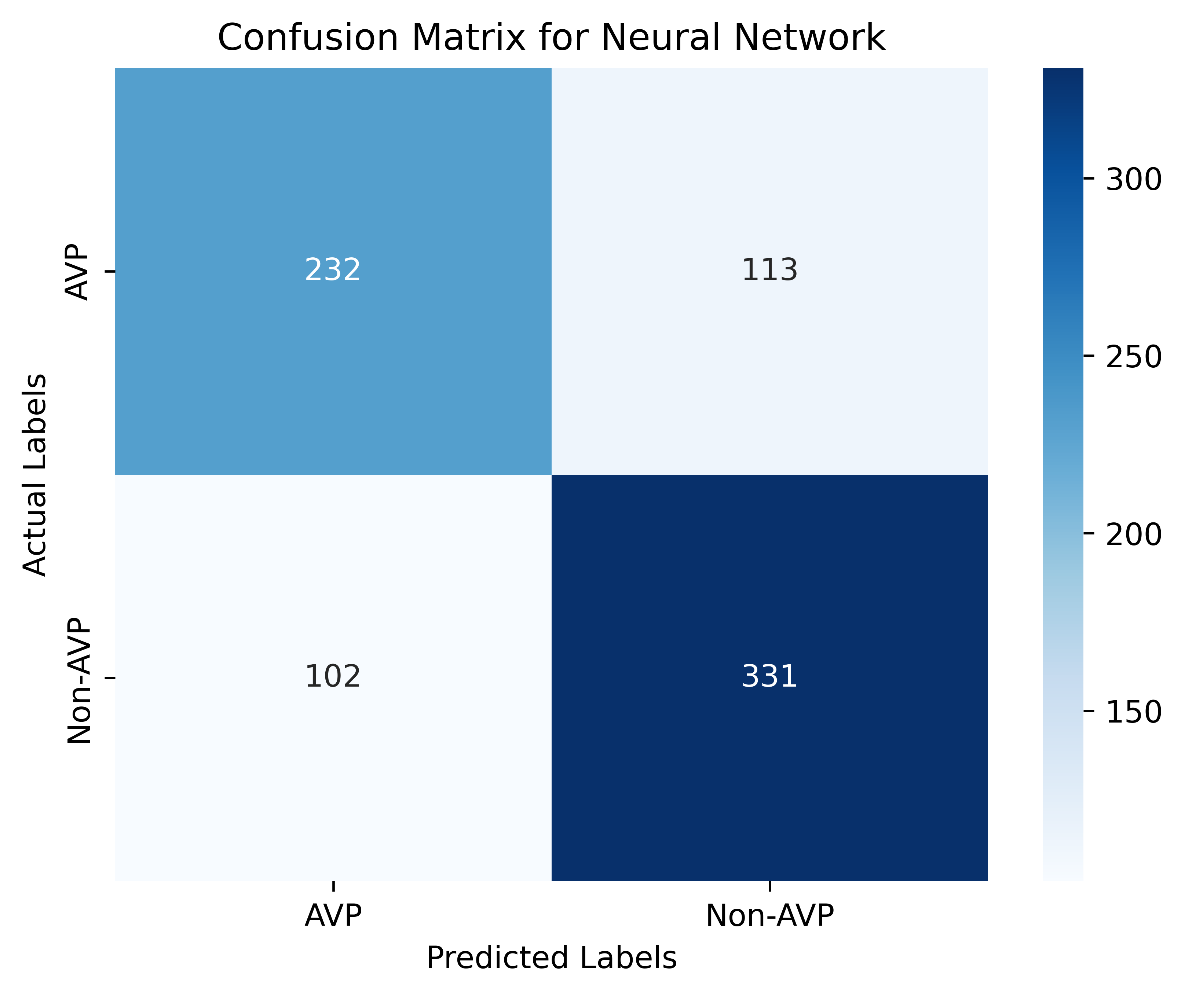

**Supplementary Figure 14: Confusion matrix of the Neural Network model.**

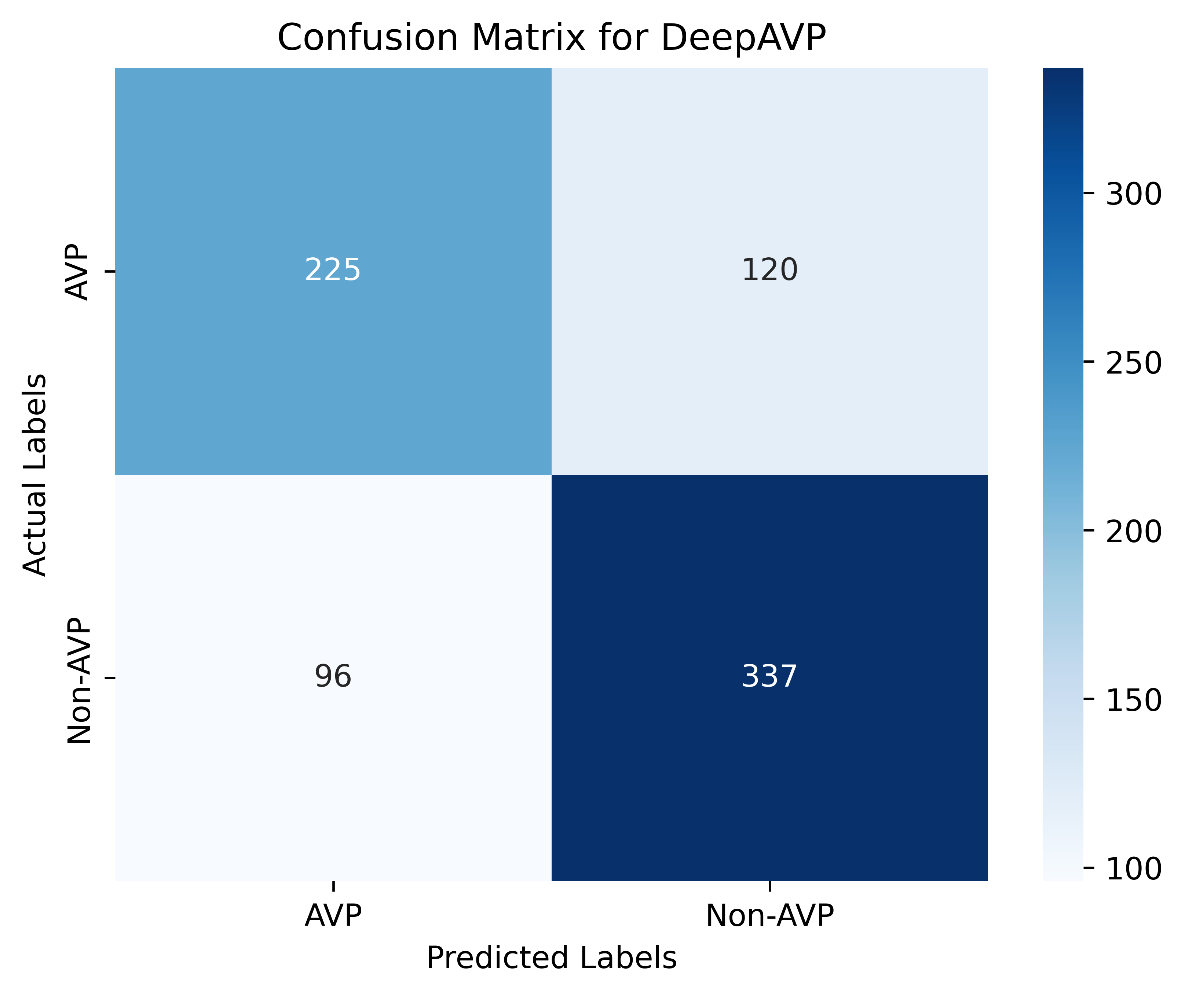

**Supplementary Figure 15: Confusion matrix of the DeepAVP model.**

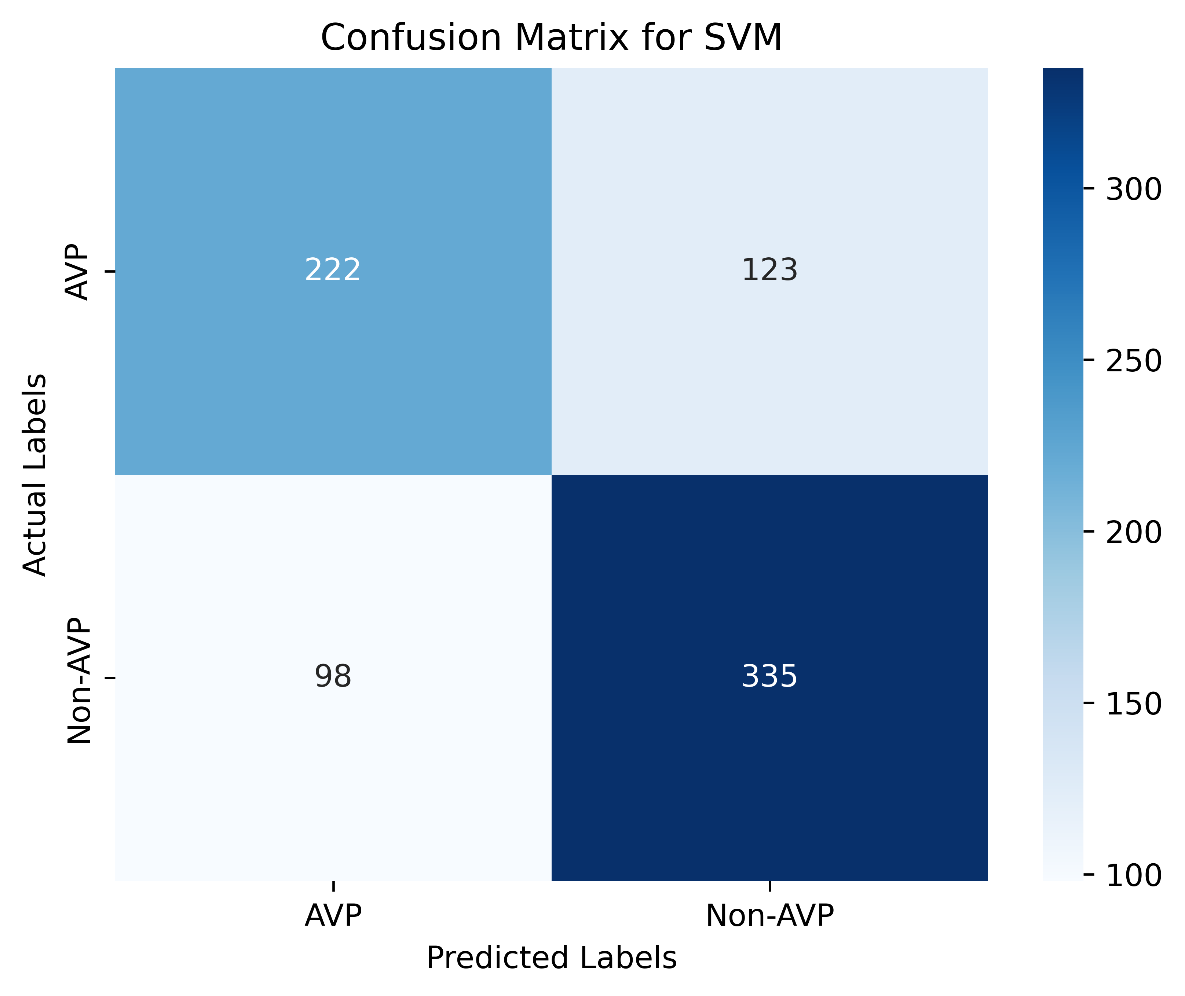

**Supplementary Figure 16: Confusion matrix of the Support Vector Machine model.**

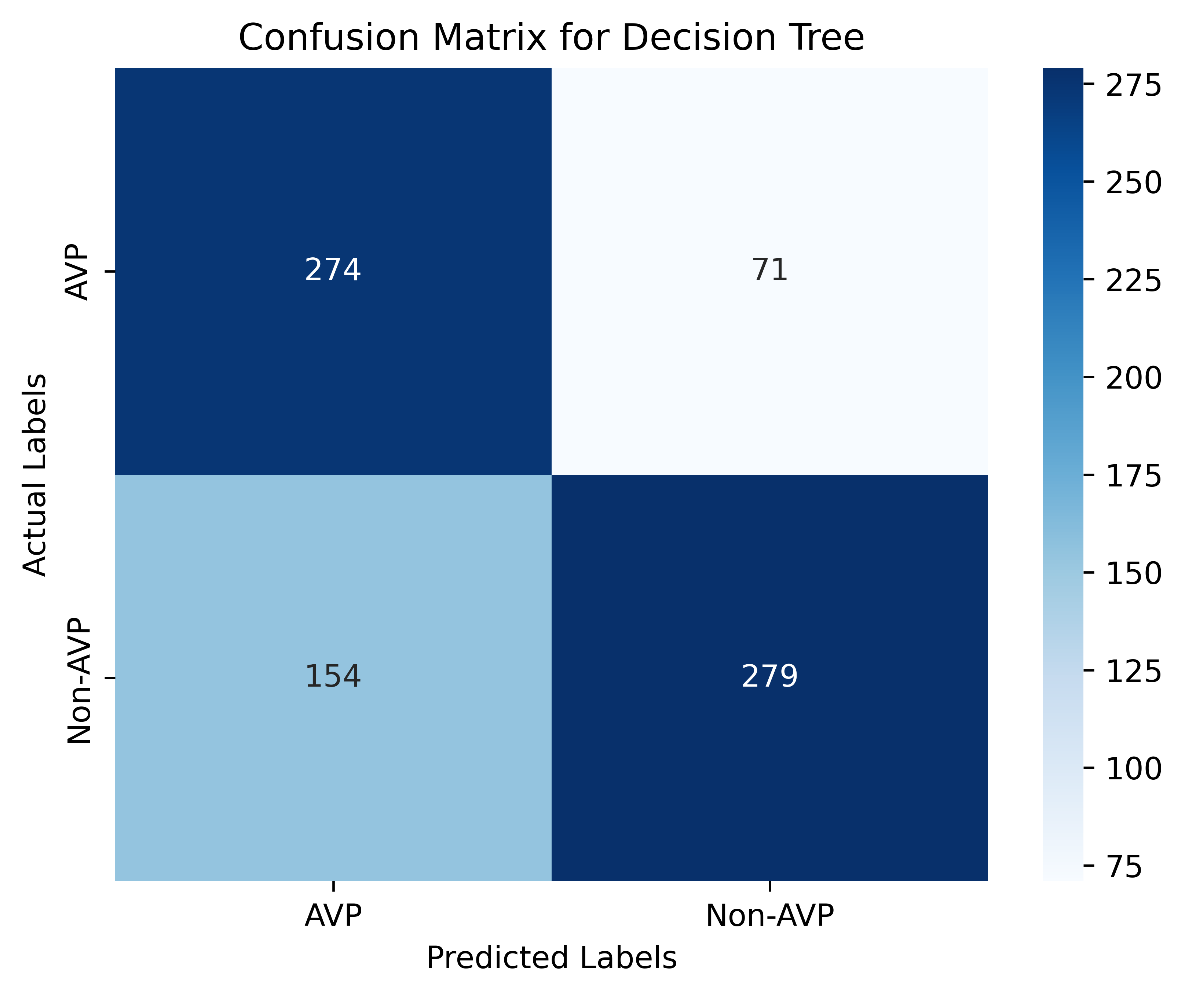

**Supplementary Figure 17: Confusion matrix of the Decision Tree model.**

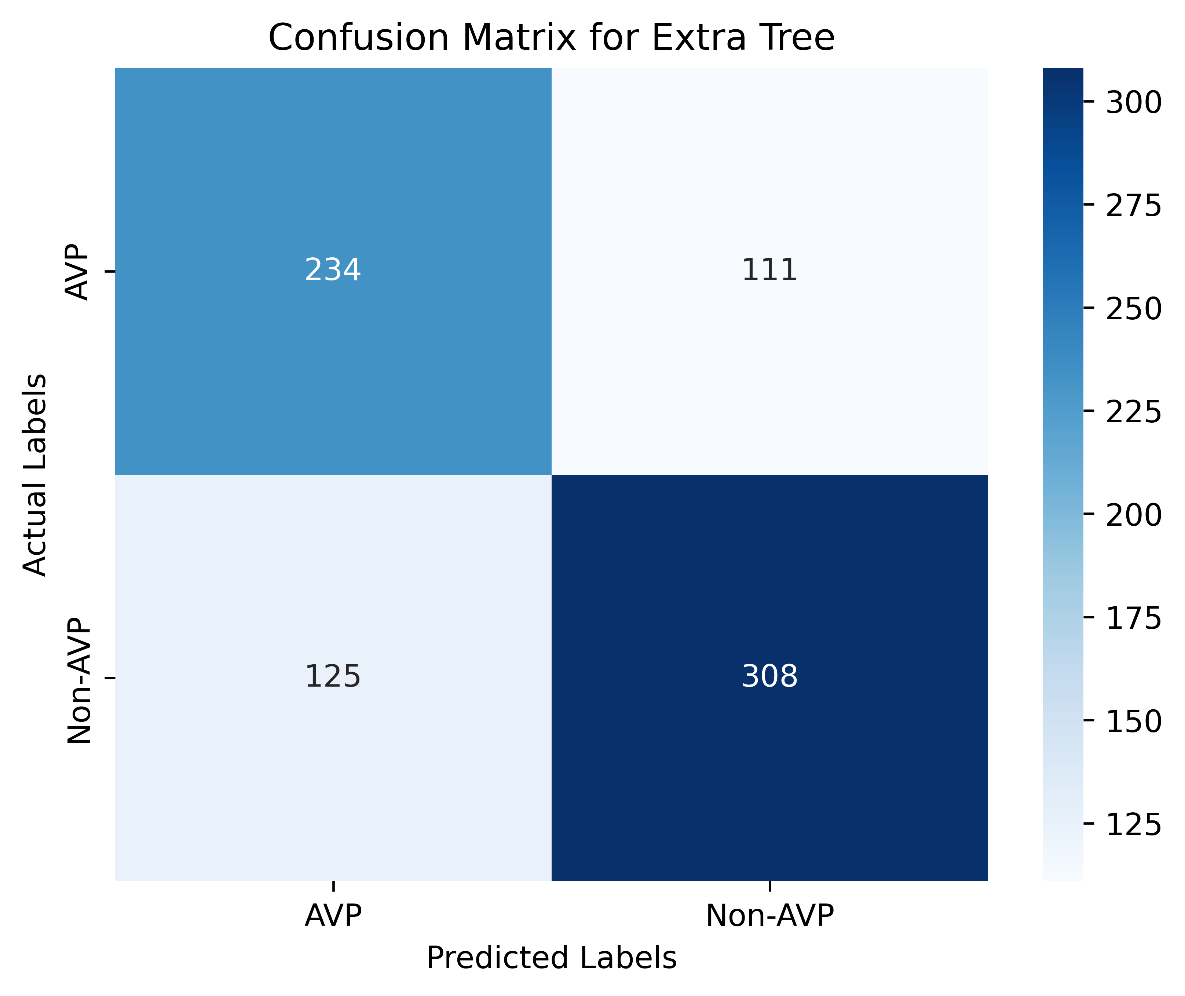

**Supplementary Figure 18: Confusion matrix of the Extra Tree model.**

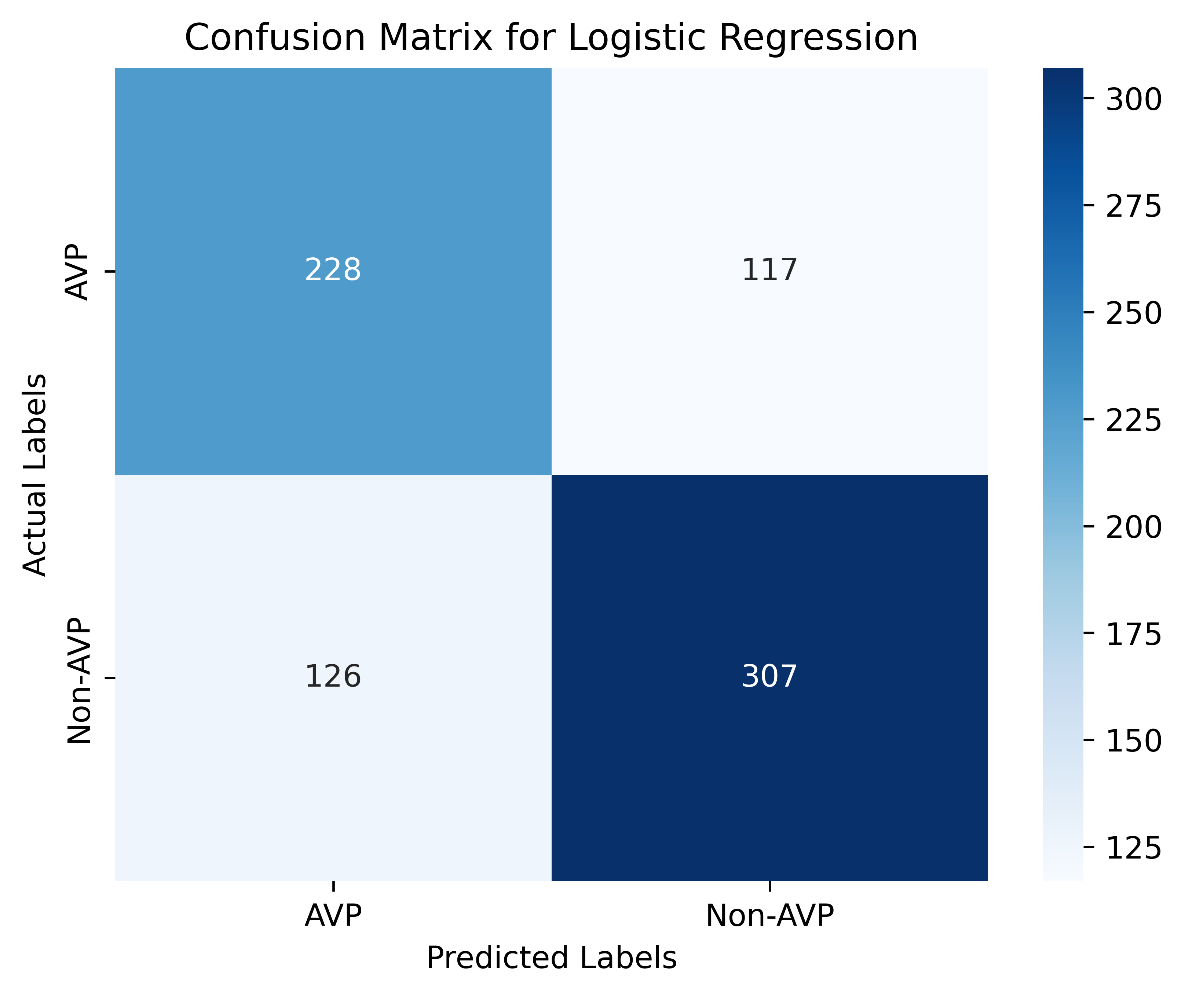

**Supplementary Figure 19: Confusion matrix of the Logistic Regression model.**

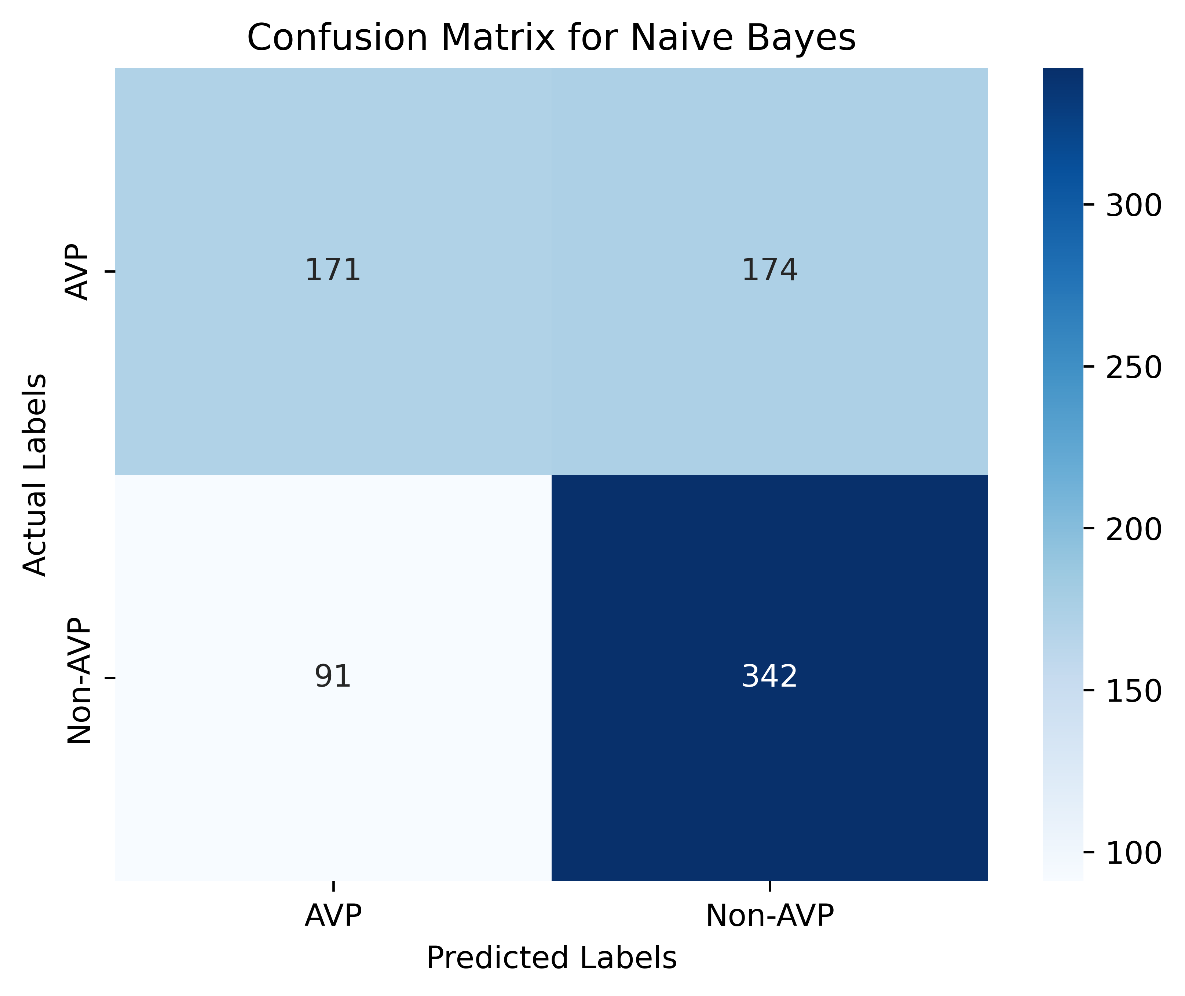

**Supplementary Figure 20: Confusion matrix of the Naive Bayes model.**

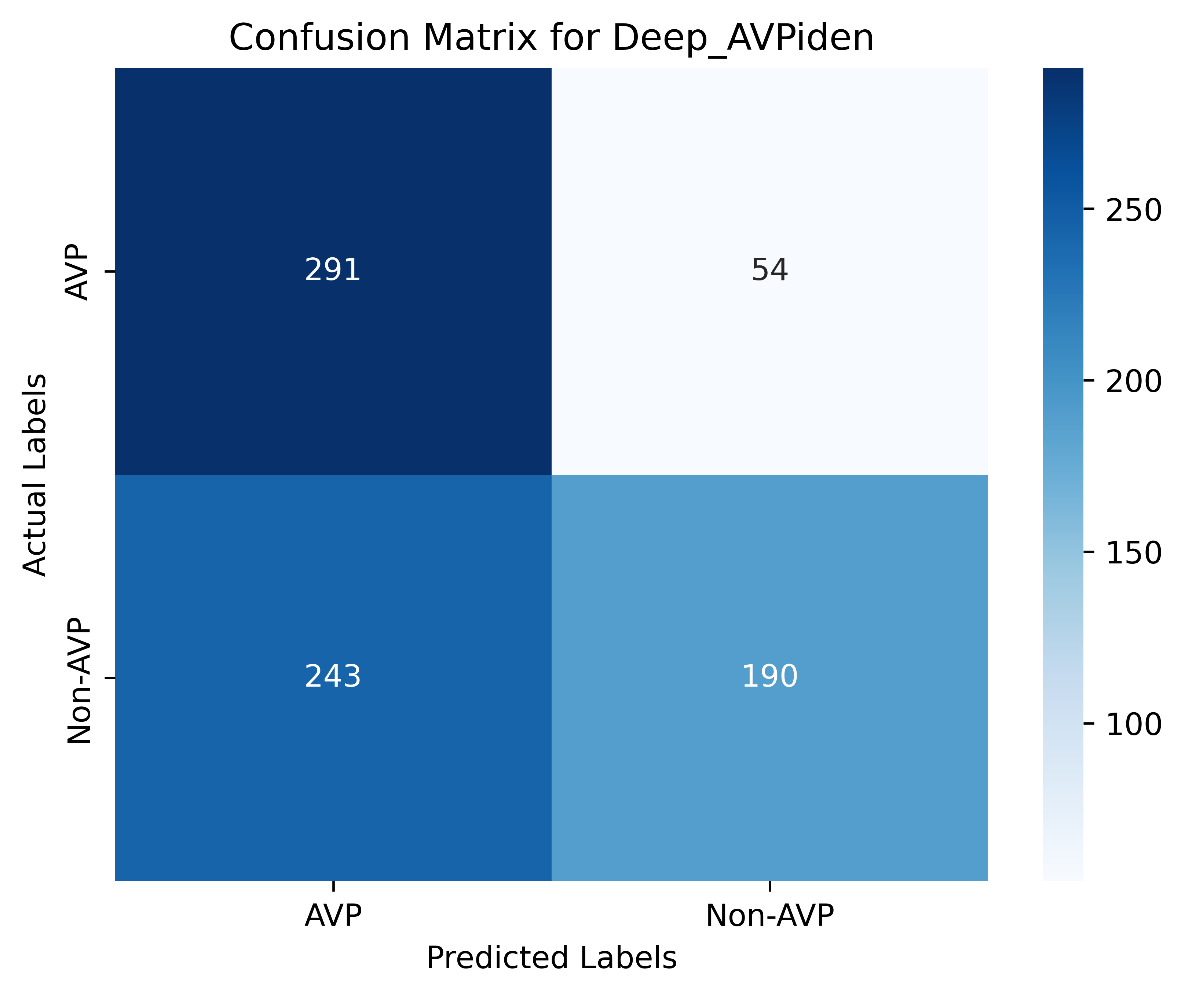

**Supplementary Figure 21: Confusion matrix of the Deep_AVPiden model.**
