## Supplementary Tables for "APDeeM: A machine Learning strategy towards Effective Peptide Vaccine Candidates Identification against Different Types of Viruses"

**Supplementary Table 1: Machine Learning Models and Hyperparameter Configuration**

| Classifier | Hyperparameters |
| --- | --- |
| Support Vector Machine | C=1.0, gamma=scale, shrinking=True, tol=1e-3, cache_size=200, max_iter=-1, decision_function_shape=ovr |
| Decision Tree | criterion=gini, splitter=best, max_depth=5, min_samples_split=2,  min_samples_leaf=1 |
| Extra Trees | n_estimators=100, max_depth=5 |
| Random Forest | n_estimators=100, max_depth=5, min_samples_split=2,  min_samples_leaf=1, random_state=42 |
| Logistic Regression | penalty=l2, tol=0.0001, C=1.0, fit_intercept=True, max_iter=100 |
| Gradient Boosting | n_estimators=100, max_depth=5, min_samples_split=2,  min_samples_leaf=1 |
| K-Nearest Neighbors | n_neighbors=5, leaf_size=30 |
| Multi-Layer Perceptron | hidden_layer_sizes=(100,), activation=relu, solver=adam,  alpha=0.0001, learning_rate_init=0.001, max_iter=200 |
| Gaussian Naive Bayes | priors=None, var_smoothing=1e-09 |
| AdaBoost | n_estimators=100 |
| XGBoost | use_label_encoder=False, eval_metric=logloss, n_estimators=100 |
| Balanced Random Forest | n_estimators=100, random_state=42 |

**Supplementary Table 2: Deep Learning Models and Hyperparameter Configuration**

| Model Name | Model Type | Hyperparameters / Layers |
| --- | --- | --- |
| DeepAVP | Hybrid LSTM-CNN Model | LSTM(64), Dropout(0.5), Conv1D(32, kernel_size=3), Dense(64, activation=relu),  Output: Dense(num_classes, activation=softmax) |
| Deep_AVPiden | Temporal Convolutional Network | Conv1D(32, kernel_size=3, dilation_rate=1), Dropout(0.3), GlobalAveragePooling1D,  Dense(64, activation=relu), Output: Dense(1, activation=sigmoid) |
| Deep_AVPpred | ResNet-like 1D CNN | Conv1D(64, kernel_size=3), BatchNormalization, MaxPooling1D,  Dense(128, activation=relu), Dropout(0.5),  Output: Dense(num_classes, activation=softmax) |
| AntiCVP_Deep | Bidirectional LSTM with Attention | Bidirectional LSTM(64), Attention Layer, GlobalAveragePooling1D, Dense(128, activation=relu),  Dropout(0.5), Output: Dense(num_classes, activation=softmax) |

**Supplementary Table 3: Ensemble Models and Hyperparameter Configuration**

| **Model Name** | **Classifier** | **Hyperparameters** |
| --- | --- | --- |
| Ensemble 1 | Gradient Boosting | n_estimators=200, learning_rate=0.1, max_depth=10, min_samples_split=2, min_samples_leaf=1, random_state=42 |
|  | Random Forest | n_estimators=200, max_depth=10, min_samples_split=2, min_samples_leaf=1, max_features=sqrt, random_state=42 |
|  | K-Nearest Neighbors | n_neighbors=5, weights=uniform, algorithm=auto, leaf_size=30, metric=minkowski |
|  | AdaBoost | n_estimators=200, learning_rate=0.1, random_state=42 |
| Ensemble 2 | Gradient Boosting | n_estimators=200, learning_rate=0.1, max_depth=10, min_samples_split=2, min_samples_leaf=1, random_state=42 |
|  | K-Nearest Neighbors | n_neighbors=5, weights=uniform, algorithm=auto, leaf_size=30, metric=minkowski |
|  | AdaBoost | n_estimators=200, learning_rate=0.1, random_state=42 |
|  | Random Forest (meta-estimator) | n_estimators=200, max_depth=10, min_samples_split=2, min_samples_leaf=1, max_features=sqrt, random_state=42 |
| Ensemble 3 | XGBoost | use_label_encoder=True, eval_metric=logloss, n_estimators=200, max_depth=10, learning_rate=0.1, subsample=0.8, colsample_bytree=0.8, gamma=0, min_child_weight=1 |
|  | Balanced Random Forest | n_estimators=200, criterion=gini, max_depth=10, min_samples_split=2, min_samples_leaf=1, bootstrap=True, class_weight=balanced, random_state=42 |
|  | K-Nearest Neighbors | n_neighbors=5, weights=uniform, algorithm=auto, leaf_size=30, metric=minkowski |
|  | AdaBoost | n_estimators=200, learning_rate=0.1, random_state=42` |
| Ensemble 4 | XGBoost | use_label_encoder=True, eval_metric=logloss, n_estimators=200, max_depth=10, learning_rate=0.1, subsample=0.8, colsample_bytree=0.8, gamma=0, min_child_weight=1` |
|  | K-Nearest Neighbors | n_neighbors=5, weights=uniform, algorithm=auto, leaf_size=30, metric=minkowski |
|  | AdaBoost | n_estimators=200, learning_rate=0.1, random_state=42` |
|  | Balanced Random Forest (meta-estimator) | n_estimators=200, criterion=gini, max_depth=10, min_samples_split=2, min_samples_leaf=1, bootstrap=True, class_weight=balanced, random_state=42 |
| Ensemble Learner | Genetic Algorithm | Population size=10, Generations=50, Mutation rate=0.1 |
|  | XGBoost | use_label_encoder=False, eval_metric=logloss, n_estimators=100 |
|  | K-Nearest Neighbors | n_neighbors=5, leaf_size=30 |
|  | Extra Trees | n_estimators=100, max_depth=5 |
|  | AdaBoost | `n_estimators=100` |

**Supplementary Table 4: Model Performance Metrics**

| Model | Accuracy  (%) | F1 Score  (%) | Recall  (%) | Precision  (%) | AUC Score  (%) | MCC  (%) | MAE  (%) | RMSE  (%) |
| --- | --- | --- | --- | --- | --- | --- | --- | --- |
| SVM | 71.59 | 75.20 | 77.37 | 73.14 | 70.86 | 42.12 | 28.40 | 53.30 |
| Decision Tree | 71.08 | 71.26 | 64.43 | 79.71 | 71.93 | 43.79 | 28.92 | 53.78 |
| Extra Tree | 69.67 | 72.30 | 71.13 | 73.51 | 69.48 | 38.82 | 30.33 | 55.08 |
| Random Forest | 74.29 | 77.53 | 79.68 | 75.49 | 73.61 | 47.64 | 25.70 | 50.70 |
| Logistic Regression | 68.77 | 71.65 | 70.90 | 72.41 | 68.49 | 36.90 | 31.23 | 55.89 |
| Gradient Boosting | 84.83 | 86.53 | 87.53 | 85.55 | 84.49 | 69.21 | 15.17 | 38.94 |
| K-Nearest Neighbors | 73.78 | 76.82 | 78.06 | 75.61 | 73.23 | 46.69 | 26.22 | 51.21 |
| Neural Network | 72.37 | 75.48 | 76.44 | 74.55 | 71.84 | 43.85 | 27.63 | 52.57 |
| Naive Bayes | 65.94 | 72.08 | 78.98 | 66.28 | 64.27 | 30.01 | 34.06 | 58.36 |
| AdaBoost | 78.02 | 79.95 | 78.75 | 81.19 | 77.93 | 55.67 | 21.98 | 46.88 |
| XGBoost | 84.96 | 86.63 | 87.53 | 85.75 | 84.63 | 69.47 | 15.03 | 38.78 |
| Balanced RF | 83.16 | 84.93 | 85.22 | 84.63 | 85.90 | 65.86 | 16.83 | 41.03 |
| **Ensemble 1** | **85.99** | **87.60** | **88.91** | **86.32** | 85.62 | **71.55** | **14.01** | **37.43** |
| Ensemble 2 | 84.06 | 85.68 | 85.68 | 85.68 | 83.86 | 67.71 | 15.94 | 39.92 |
| Ensemble 3 | 83.29 | 85.23 | 86.61 | 83.89 | 82.87 | 66.05 | 16.71 | 40.88 |
| Ensemble 4 | 82.26 | 83.99 | 83.60 | 84.38 | 82.09 | 64.11 | 17.74 | 42.12 |
| Ensemble Learner | 79.05 | 80.76 | 78.98 | 82.61 | **88.30** | 57.86 | 20.95 | 45.77 |
| DeepAVP | 72.24 | 72.11 | 72.24 | 72.12 | 79.40 | 43.44 | 27.76 | 52.96 |
| Deep_AVPiden | 61.83 | 56.13 | 43.88 | 77.87 | 73.97 | 30.22 | 38.17 | 61.79 |
| Deep_AVPpred | 75.58 | 74.77 | 75.58 | 76.71 | 83.30 | 50.89 | 24.42 | 49.42 |
| AntiCVP_Deep | 78.02 | 78.04 | 78.02 | 78.06 | 86.59 | 55.54 | 21.98 | 46.88 |
